# Economic and Social Modulations of Innate Decision-Making in Mice Exposed to Visual Threats

**DOI:** 10.1101/2025.05.12.653401

**Authors:** Zhe Li, Jiahui Wang, Yidan Sun, Jialin Li, Ling-yun Li, Ya-tang Li

## Abstract

When confronted by predators, animals make innate decisions with rapid reaction times—a trait shaped by natural selection to maximize survival. However, rapid reactions are effective only when grounded in accurate judgments and appropriate choices, which often require cognitive control. To address how such choices are shaped, we developed a behavioral paradigm to investigate how threat intensity, reward value, and social hierarchy shape decision-making in foraging mice exposed to overhead visual threats. Using a machine learning-based approach, we classified defensive responses into four distinct decision types. Mice showed rapid habituation to repeated looming threats, with substantial inter-individual variability in the rate of habituation. Across both early and late phases of habituation, threat intensity emerged as the primary determinant of decision-making, strongly biasing behavior toward escape. In contrast, the influence of reward value was context-dependent and became evident primarily in the late phase: under low-threat conditions, higher reward value suppressed defensive responses, consistent with value-based decision theory; whereas under high-threat conditions, higher reward value promoted escape, potentially reflecting heightened vigilance. Innate decision-making was further modulated by social hierarchy, with dominant mice showing greater vigilance and a stronger bias toward risk-averse behaviors, while subordinates were more reward-driven. To understand the underlying decision-making process, we developed a drift-diffusion leaky integrator model that successfully captures how threat intensity, reward value, and vigilance interact to shape defensive decisions. Together, these findings reveal how economic and social factors modulate innate decisions and provide a computational framework for understanding the interplay between instinctive reactions and cognitive control.

## 1 Introduction

How animals make decisions among alternative actions is a key question in neuroscience. In natural environments, decisions arise from the integration of sensory inputs, internal states, and learned experience (Meister, 2022). In the laboratory, both learned and innate paradigms have been used to study decision-making. Some learned paradigms, such as two-alternative forced choice (2-AFC) and Go/No-go tasks (Andermann et al., 2010; Burgess et al., 2017), often require weeks of training in head-fixed animals; whereas other learned paradigms require less training over several days in freely moving animals, including maze exploration (Barnes, 1979; Morris, 1984; Rosenberg et al., 2021; Small, 1901), foraging decisions (Hayden, 2018; Hayden et al., 2011; Steiner and Redish, 2014), and active avoidance (Bravo-Rivera et al., 2014; Moscarello and LeDoux, 2013; Mowrer and Lamoreaux, 1946). While these paradigms have significantly advanced our understanding of the neural mechanisms underlying learned decisions, the training itself may engage neural circuits that differ from those evolved for innate decision-making (Steinmetz et al., 2019).

In contrast, innate decisions—such as whether to escape from an approaching aerial predator (De Franceschi et al., 2016; Yilmaz and Meister, 2013) or a simulated ground threat like a robogator (Amir et al., 2015; Choi and Kim, 2010)—are executed without prior training and may more directly reflect the function of neural circuits shaped by natural selection. In complex environments, making correct and rapid innate defensive decisions often requires cognitive control to assess risk and evaluate alternative defensive strategies (Evans et al., 2019). Defensive responses to terrestrial predators depend on predator–prey distance (Ydenberg and Dill, 1986) and are well described by the predatory imminence continuum theory (Fanselow and Lester, 1988). Responses to approaching aerial predators are likely scaled similarly according to the perceived threat intensity, which is influenced by the physical properties of the threat (De Franceschi et al., 2016; Liden and Herberholz, 2008; Tammero and Dickinson, 2002; Yang et al., 2020; Yilmaz and Meister, 2013), prior experience (Vale et al., 2017), and environmental context, including housing conditions (Lenzi et al., 2022). At the neural level, the superior colliculus (SC) serves as a central hub for processing looming-evoked defensive responses, with distinct output pathways mediating escape versus freezing behaviors (Evans et al., 2018; Shang et al., 2018; Wei et al., 2015; Zhou et al., 2019).

Despite this progress, important gaps remain. Mice are social animals, and most predator encounters occur during foraging, yet it is unclear how they weigh perceived threat and reward when making defensive decisions, or how these decisions are influenced by social hierarchy. To address these questions, we developed an ecologically relevant behavioral paradigm to investigate decision-making in foraging mice exposed to overhead visual threats. Our results demonstrate that defensive choices are jointly influenced by threat intensity, reward value, and social hierarchy. Furthermore, we identify vigilance as a key factor linking threat and reward information to behavioral choice. These findings establish a behavioral framework for future investigations into the neural mechanisms underlying the cognitive control of innate decision-making.

## 2 Results

### 2.1 A behavioral paradigm for studying innate decision-making in mice

To investigate how animals make innate decisions in natural environments, we designed a behavioral paradigm to simulate the defensive behavior of foraging mice in the wild. In this paradigm, a group of 2–5 co-housed mice was placed in a nest, with each individual identified using a radio frequency identification (RFID) tag. Only one mouse was allowed to enter a linear arena to receive the reward delivered at the end. As the mouse approached the reward, an overhead expanding dark disc that mimics the approach of an aerial predator was triggered (Figure 1A), forcing the animal to decide whether to risk getting the reward or to seek safety in the nest. Behavioral data were recorded using a ground-mounted camera, and DeepLabCut was used to track the movements of the mouse’s nose and tail base (Mathis et al., 2018) (Figure S1A). Compared with the exploratory phase, looming stimuli significantly increased the frequency of arena entries but decreased the duration of each visit (Figures 1B–C).

**Figure 1:**
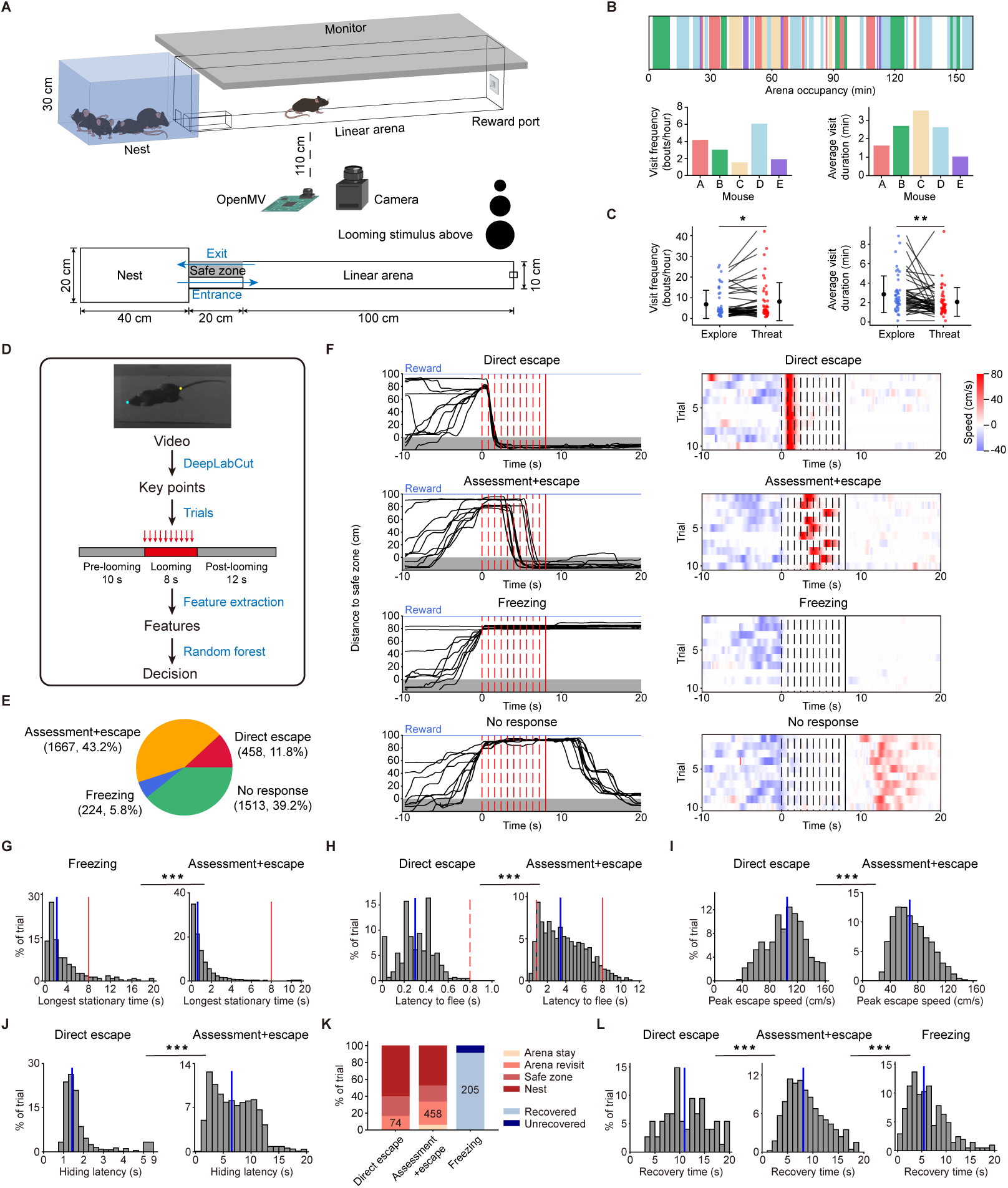
A behavioral paradigm for investigating innate decision-making in mice. (A) Schematic of the behavioral assay (3D and top-down views). (B) Top: arena occupancy patterns for five example mice in one session. Bottom: visit frequency and average duration for each visit. Different colors mark the mouse’s identity. (C) Visit frequency and duration under exploration and threat conditions. Error bars represent standard deviation. *n* = 46 mice; paired *t*-test. (D) Pipeline for behavioral classification. (E) Distribution of decisions across 3862 trials from 140 mice. (F) Left: distance to the safe zone over time for four decision types (10 example trials each). Red dashed lines mark the onset of each stimulus repetition; solid lines mark the end of the last repetition. Gray shading indicates the safe zone. Right: locomotion speed toward the safe zone over time for the same trials. Positive speed indicates movement toward the safe zone. (G) Distribution of the longest stationary time for “freezing” and “assessment+escape”. *n* = 224 and 1667 trials, Mann–Whitney *U* test. Blue lines mark the median; red solid lines mark the end of the last repetition. (H) Distribution of latency to flee for “direct escape” and “assessment+escape”. *n* = 458 and 1667 trials, Mann–Whitney *U* test. Blue lines mark the median; red dashed and solid lines mark the end of the first and last repetitions, respectively. (I) Distribution of peak speed for “direct escape” and “assessment+escape”. *n* = 458 and 1667 trials, Mann–Whitney *U* test. (J) Distribution of hiding latency in the safe zone for “direct escape” and “assessment+escape”. *n* = 455 and 1563, Mann–Whitney *U* test. (K) Distribution of post-decision behavioral states. Numbers indicate the trial counts in which mice recovered from the fear state within 20 s after stimulus onset. (L) Distribution of fear recovery time for “direct escape”, “assessment+escape”, and “freezing”. *n* = 74, 458, and 205, Kruskal–Wallis test followed by Dunn’s *post hoc* test with Holm correction. For all panels: \**P <* 0.05, \*\**P <* 0.01, \*\*\**P <* 0.001.

To identify distinct behavioral patterns in response to looming stimuli, we defined 19 behavioral features from key body points of the mouse and fed them into a random forest classifier that achieved 95% classification accuracy (Figures 1D and S1B–C, see Materials and Methods). Across 3862 trials, decisions were categorized into four types: direct escape (11.8%, Video 1), escape after assessment (assessment+escape, 43.2%, Video 2), freezing (5.8%, Video 3), and no response (39.2%, Video 4) (Figures 1E–F). A key distinction between assessment+escape and freezing decisions was the duration of stationary behavior: mice that decided to freeze remained stationary significantly longer than those in the assessment+escape group (Figure 1G). Furthermore, the latency to flee differed significantly between direct escape and assessment+escape decisions (Figure 1H).

We hypothesized that fear level increases progressively from freezing to assessment+escape to direct escape. This hypothesis is supported by the observations that mice in the direct escape group exhibited higher escape speeds (Figure 1I) and shorter hiding latencies (Figure 1J). Additionally, the proportion of mice that recovered within 20 s after stimulus onset was about 16.2%, 27.5%, and 91.5% for direct escape, assessment+escape, and freezing, respectively (Figure 1K). Consistently, recovery time decreased progressively across these decision types (Figure 1L). Our findings align with the predatory imminence continuum theory (Fanselow and Lester, 1988), which proposes that defensive responses become more intense as the perceived threat intensity increases.

### 2.2 Mice make economic decisions modulated by vigilance

In the wild, most prey–predator encounters occur during foraging, and prey have evolved to maintain high vigilance to enhance survival. Yet how food value, threat intensity, and vigilance interact to shape defensive decisions remains unclear. To address this question, we developed a behavioral assay that simulates how a foraging animal responds to an approaching aerial predator under different threat and reward conditions (Figure 2A). To maintain a consistent internal state across conditions, mice were not water-deprived. The higher reward value of sucrose over water was validated by measuring consumption during exploration (Figure S3A).

**Figure 2:**
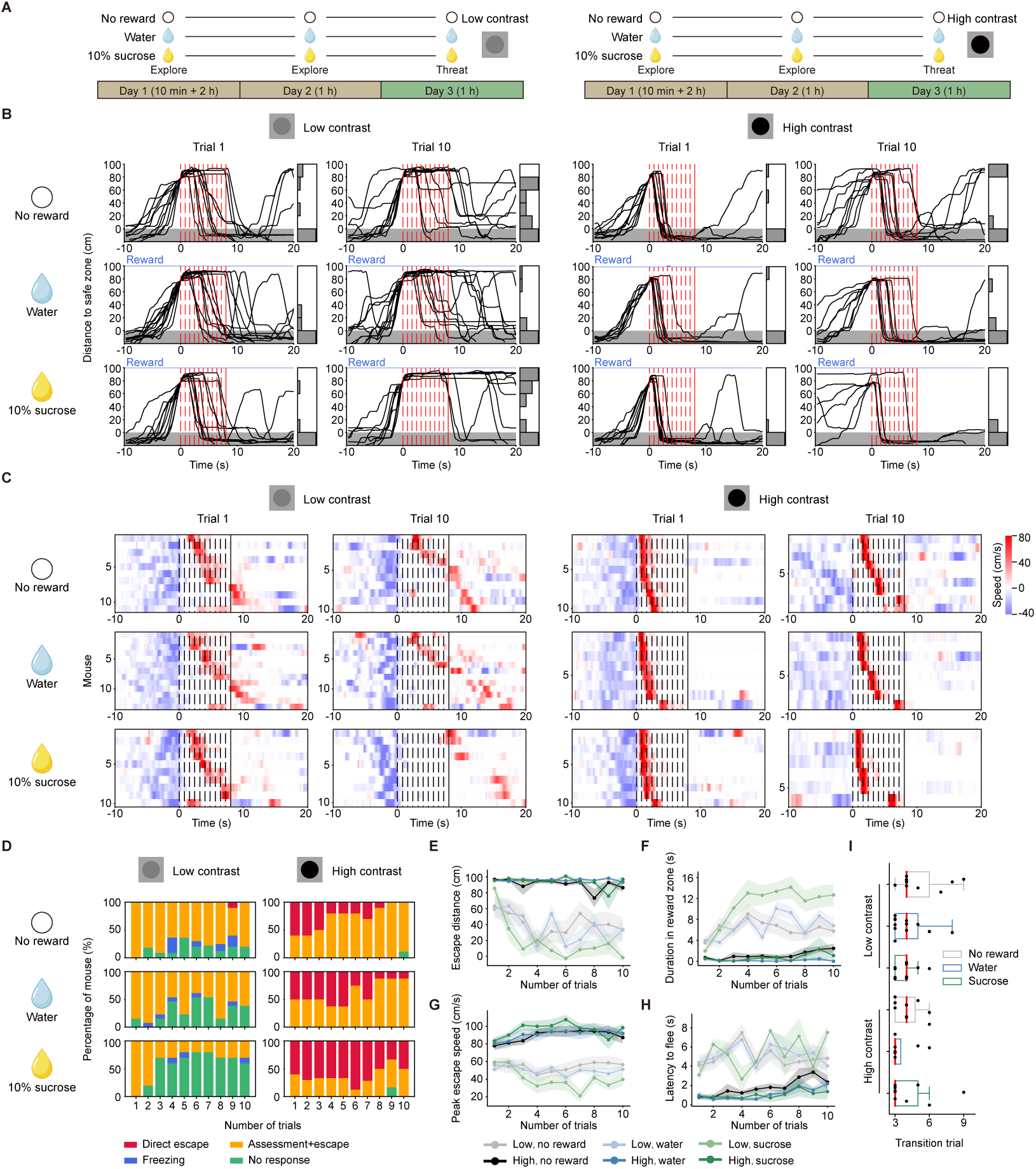
Rapid learning shapes economic decisions under threat. (A) Schematic of the behavioral assay for studying the economic modulation of innate decision-making. (B-C) Distance to the safe zone and locomotion speed over time for the first and tenth trials in response to low- and high-contrast looming stimuli across different reward conditions. The right inset in (B) shows the distribution of distance to the safe zone at the end of each trial. Dashed lines mark the start of each stimulus, and solid lines mark the stimulus offset. *n* = 11 (no reward, low), 13 (water, low), 10 (sucrose, low), 10 (no reward, high), 8 (water, high), 10 (sucrose, high) mice. (D) Summary of the decisions across the first 10 trials under different threat and reward conditions. *n* = 11, 13, 10, 10, 8, 10 mice. (E–H) Escape distance under threat, duration in the reward zone, peak escape speed, and latency to flee across trials in all conditions. Shading denotes the standard error of the mean. (I) Transition trials marking the shift between phases across conditions. Red lines indicate the median; boxes span the interquartile range (IQR); whiskers extend to 1.5 *×* IQR beyond the box.

During the first trial, mice showed distinct trajectories and speed profiles across threat and reward conditions (Figures 2B–C). These patterns changed rapidly within the first 10 trials, indicating fast habituation to the looming stimulus. This learning was reflected in altered decision patterns (Figure 2D) and changes in escape distance, time spent in the reward zone, peak escape speed, and latency to flee (Figures 2E–H). To account for both learning and individual variability, we segmented trials for each mouse into early and late phases using a change-point detection approach on the learning curves (Figures 2I and S3C–D, see Materials and Methods).

In the early phase, behavior was shaped predominantly by the level of threat. Under higher threat, mice were more likely to choose direct escape decisions (Figure 3A), with longer escape distances (Figure 3B) and less time spent in the reward zone (Figure 3C), suggesting a trade-off between threat avoidance and reward pursuit. Notably, latency to flee decreased with higher threat (Figure 3D), consistent with heightened vigilance (Buck, 1966). This threat-dependent change in vigilance was further supported by the longer interval before re-entering the trigger zone and slower foraging speed (Figures 3E–F and S4), consistent with findings in birds showing that elevated predation risk increases vigilance and reduces feeding time (Caraco et al., 1980). In addition, escape speed increased significantly with threat (Figure 3G), reflecting the combined effects of perceived higher risk and heightened vigilance. Analyses restricted to the first trial yielded consistent results (Figure S5).

**Figure 3:**
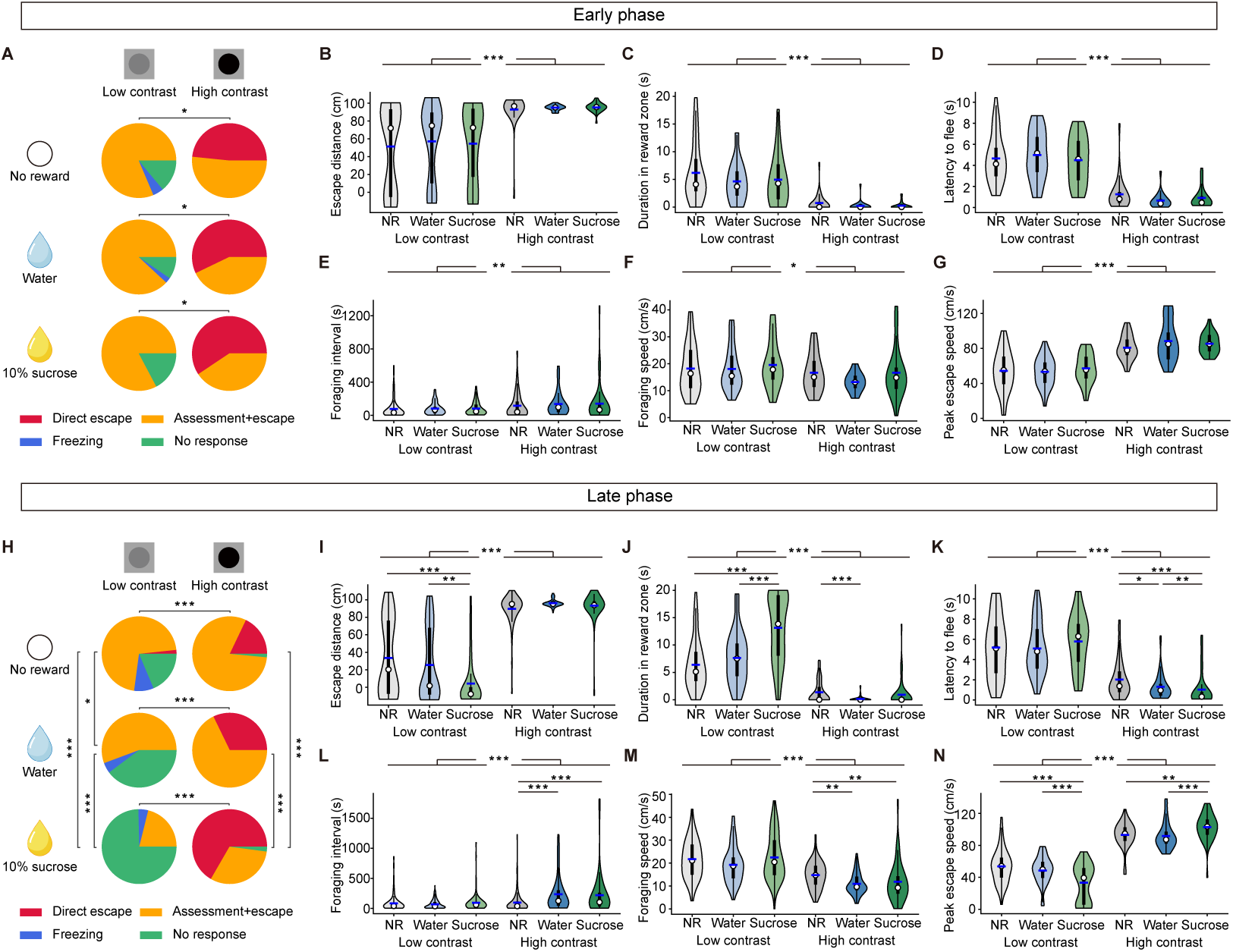
Mice make economic decisions modulated by vigilance. (A) Distribution of decisions in the early phase under six experimental conditions. *n* = 43 trials from 11 mice (no reward, low), 42 trials from 13 mice (water, low), 29 trials from 10 mice (sucrose, low), 31 trials from 10 mice (no reward, high), 21 trials from 8 mice (water, high) and 32 trials from 10 mice (sucrose, high); chi-squared test. (B–G) Escape distance under threat, duration in the reward zone, latency to flee, foraging interval, foraging speed, and peak escape speed in the early phase across all conditions. *n* = 43, 42, 29, 31, 21, 32 trials for B, C, F, G; *n* = 35, 37, 24, 31, 21, 32 trials for D; *n* = 116, 84, 54, 58, 36, 48 intervals for E; Scheirer–Ray–Hare test with *post hoc* Dunn’s test (Holm correction). (H) Distribution of decisions in the late phase under six experimental conditions. *n* = 59 trials from 11 mice (no reward, low), 88 trials from 13 mice (water, low), 71 trials from 10 mice (sucrose, low), 67 trials from 10 mice (no reward, high), 59 trials from 8 mice (water, high) and 48 trials from 9 mice (sucrose, high); chi-squared test. (I-N) Escape distance under threat, duration in the reward zone, latency to flee, foraging interval, foraging speed, and peak escape speed in the late phase across all conditions. *n* = 59, 88, 71, 67, 59, 48 trials for I, J, M, N; *n* = 43, 49, 15, 66, 59, 47 trials for K; *n* = 116, 182, 114, 137, 71, 57 intervals for L; Scheirer–Ray–Hare test with *post hoc* Dunn’s test (Holm correction). For all panels: \**P <* 0.05, \*\**P <* 0.01, \*\*\**P <* 0.001.

In the late phase, both threat intensity and reward value contributed to behavior. Threat remained a dominant factor, while reward exerted a significant, context-dependent influence. Under low-threat conditions, decisions were primarily driven by perceived reward value. As reward value increased, mice exhibited fewer defensive responses (Figure 3H), indicating that they weighed threat and reward to make economic decisions. Higher reward values led to shorter escape distance (Figure 3I) and longer stays in the reward zone (Figure 3J), supporting value-based decision-making strategies. In contrast, vigilance-related metrics, including latency to flee, foraging interval, and foraging speed, were largely unchanged across reward conditions (Figures 3K–M), suggesting that vigilance remained relatively stable. Consistently, escape speed decreased as reward value increased (Figure 3N). The value-based decision-making under low-threat conditions was further validated using within-subject comparisons to control for individual variability (Figure S6).

Under high-threat conditions, however, increasing reward value produced the opposite effect: mice showed more direct escape behaviors (Figure 3H). This shift in decision patterns was accompanied by behavioral signatures of heightened vigilance, including shorter latencies to flee, longer foraging intervals, and slower foraging speeds (Figures 3K–M). These results point to a counterintuitive mechanism: under high threat, reward value does not directly promote continued foraging but instead shapes decisions by enhancing vigilance, which in turn biases animals toward rapid escape responses. Consistent with this interpretation, the duration spent in the reward zone did not increase with reward value (Figure 3J). Collectively, these findings reveal how innate decision-making in response to looming stimuli is shaped by the dynamic interplay between perceived threat intensity and reward value, with vigilance acting as a key modulating factor.

### 2.3 Influence of social hierarchy on decision-making under threat

Mice are social animals that live in groups, where social hierarchy often plays a critical role in shaping behavior. To investigate how social rank influences decision-making under threat, we compared the responses of dominant and subordinate mice to looming stimuli. Rank within each mouse pair was determined using the tube test before and after the behavioral experiments (Figure 4A, see Materials and Methods), and only pairs with consistent rankings were included in the analysis.

**Figure 4:**
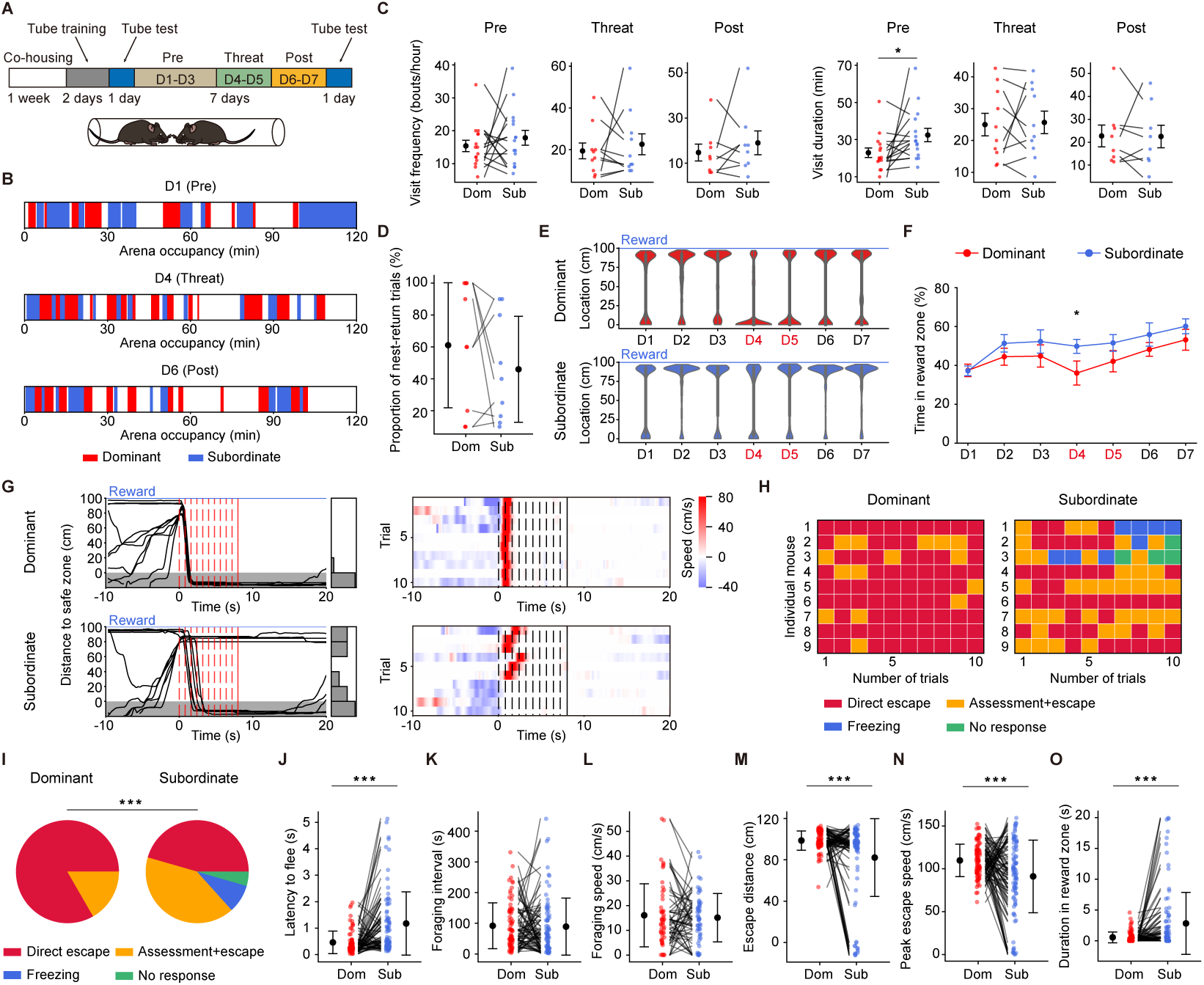
Influence of social hierarchy on innate decision-making. (A) Schematic of the behavioral assay for studying the social modulation of innate decision-making. Top, experimental timeline. Each session lasted 2 hours per mouse pair during the pre-threat, threat, and post-threat phases. Bottom, schematic of the tube test. (B) Arena occupancy for an example pair of mice during the three sessions. (C) Visit frequency and total visit duration for dominant and subordinate mice during the pre-threat, threat, and post-threat sessions. *n* = 5 pairs (pre), 5 pairs (threat), 4 pairs (post), respectively; paired *t*-test. (D) Proportion of nest-return trials in escape trials for dominant and subordinate mice. *n* = 9 pairs; paired *t*-test. (E) Distance to the safe zone over seven days for an example pair. Looming stimuli were presented on days 4 and 5. (F) Percentage of time spent in the reward zone across days. Error bars represent SEM. *n* = 9 pairs; paired *t*-test. (G) Distance to the safe zone (left) and locomotion speed (right) for an example pair. (H) Behavioral decisions across the first 10 trials for 9 mouse pairs. (I) Pie charts showing decision distributions for dominant and subordinate mice. *n* = 90, 90 trials; Stuart-Maxwell test. (J–O) Latency to flee, foraging interval, foraging speed, escape distance under threat, peak escape speed, and duration in the reward zone for dominant and subordinate mice. *n* = 78 trials for J; *n* = 75 trials for K, 53 for L, and 90 for M–O; paired *t*-test. For all panels: \**P <* 0.05, \*\**P <* 0.01, \*\*\**P <* 0.001.

We first verified that behavioral responses to the visual threat were not confounded by the presence of a social partner. Before looming exposure, subordinate mice spent more time exploring the arena than dominant mice (Figures 4B–C), which may reflect social avoidance. However, this difference disappeared during and after looming exposure, suggesting that the looming stimulus became the primary driver of behavior. To confirm this, we examined the probability of mice fleeing directly to the nest. If subordinates were avoiding dominants, they would preferentially escape to the safe zone rather than return to the nest occupied by the dominant mouse. Contrary to this prediction, dominant and subordinate mice showed similar probabilities of fleeing directly to the nest (Figure 4D), confirming that defensive responses were driven by visual threat rather than social avoidance.

When the threat and reward co-localized, only dominant mice reduced their relative time spent in the reward zone during the 2-h session (Figures 4E–F), indicating greater vigilance and risk aversion compared to subordinates. This heightened vigilance in dominant mice was further supported by reduced habituation to the looming threat (Figures 4G–H and S7), a higher proportion of direct escape decisions (Figure 4I), and shorter latencies to flee (Figure 4J). No rank differences were observed in foraging intervals or foraging speeds (Figures 4K–L). This may stem from the experimental design, in which only one mouse was allowed in the arena at a time, potentially limiting the sensitivity of these measures to detect rank-dependent differences in vigilance.

Furthermore, dominant mice prioritized threat avoidance over reward, fleeing longer distances at higher escape speeds and spending less time in the reward zone (Figures 4M–O). To rule out the possibility that the tube test itself influenced defensive behavior, we conducted additional experiments in which looming sessions occurred both before and after the tube test (Figure S8); the results remained largely consistent. Together, these findings indicate that social hierarchy biases the trade-off between reward seeking and threat avoidance, with dominant mice exhibiting heightened threat vigilance and subordinate mice showing stronger reward-driven behavior.

### 2.4 A mathematical model for innate decision-making under threat

Our experimental results demonstrate that innate decision-making in response to visual threats is influenced by perceived threat intensity, reward value, and vigilance. To understand the underlying decision-making process, we developed a mathematical model in which the evidence for escape is accumulated by a drift-diffusion leaky integrator. An escape response is triggered when the evidence level crosses a predefined threshold:

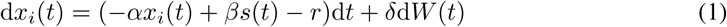

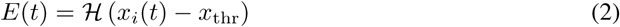

Here, *x_i_*(*t*) is the accumulated escape evidence at time *t* on the *i*th trial. The leakage rate *α* drives the evidence toward zero and is equal to the reciprocal of the integration time constant. The stimulus function *s*(*t*) denotes the normalized diameter of the looming stimulus (Figure S9A). Together with the threat gain *β*, which reflects both stimulus contrast and the animal’s vigilance level, these terms determine the perceived threat intensity and drive evidence accumulation toward escape. The term *r* represents the perceived reward value, which suppresses the accumulation of escape evidence. The stochastic term d*W* (*t*) denotes increments of a Wiener process, whose increments satisfy *W* (*t* + Δ*t*) − *W* (*t*) ∼ *N* (0, Δ*t*), and *δ* is the diffusion rate. The binary variable *E*(*t*) indicates whether an escape decision has been reached, with *H* denoting the Heaviside step function and *x*_thr_ the decision threshold. If a decision is reached within the first expansion of the looming disc, it is classified as a direct escape; otherwise, it is classified as an escape after assessment.

Model parameters are estimated using behavioral data from the late phase of the reward–threat experiments in two steps (see Materials and Methods). First, we fit the model to the data under no-reward condition to obtain the optimal leakage rate (*α* = 1.78), threat gain (low threat: *β* = 1; high threat: *β* = 1.44), diffusion rate (*δ* = 5.6), and decision threshold (*x*_thr_ = 0.71) (Figure S9B). These parameters reveal distinct decision-making dynamics under low- and high-threat conditions. Under low threat, the average evidence level remains below the escape threshold, and escape decisions emerge from stochastic fluctuations in individual trials (Figure 5A). Under high threat, the average evidence level crosses the threshold, indicating that escape decisions are mainly driven by threat gain (Figure 5B). Notably, threat gain and diffusion rate exert distinct effects on latency to flee: threat gain shifts the mean latency, while diffusion rate affects its variance. Thus, the model captures not only the observed decision patterns but also the distributions of latency to flee.

**Figure 5:**
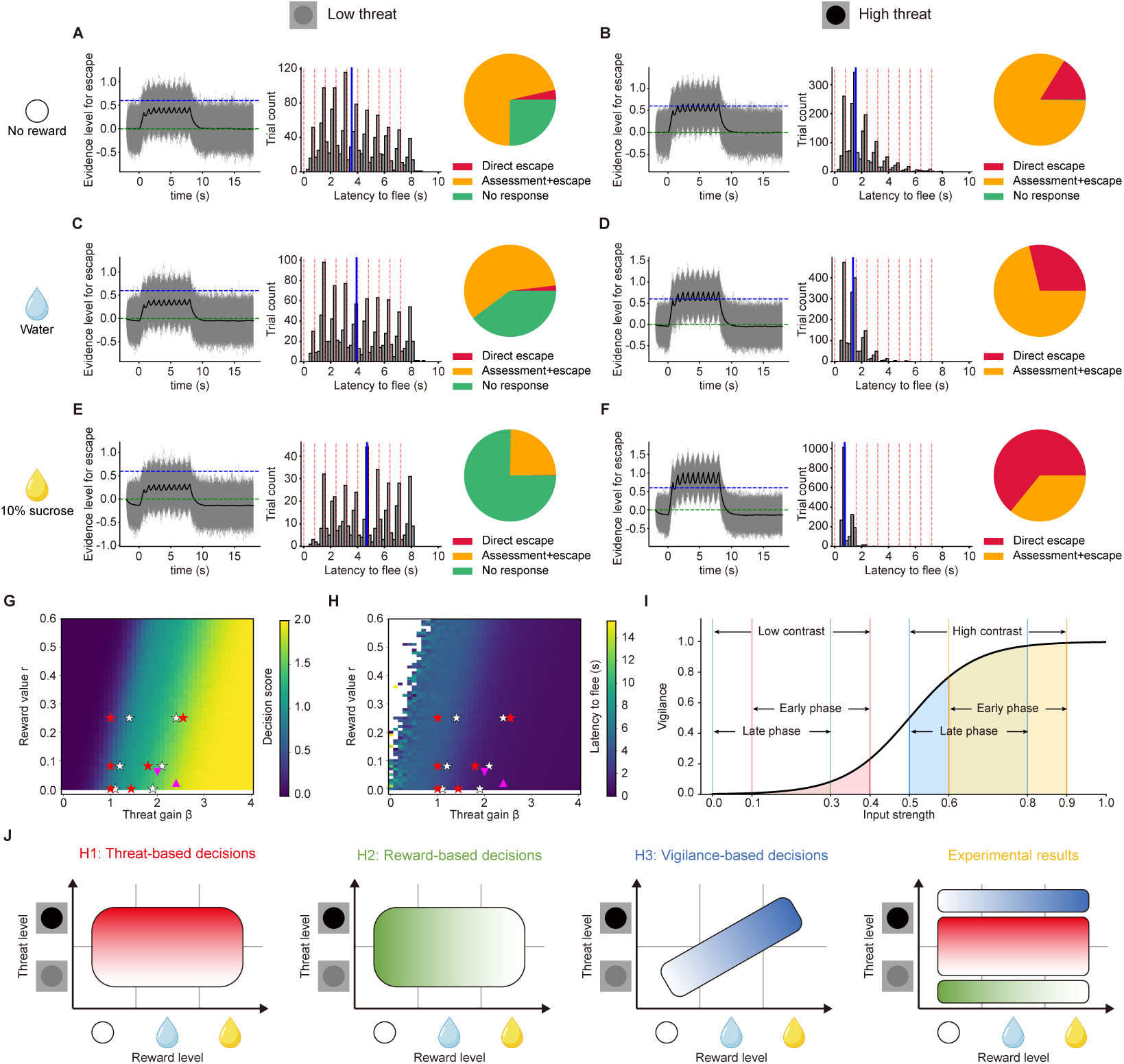
Drift-diffusion leaky integrator model for escape decisions. (A-F) Simulated accumulation of escape evidence, along with predicted latencies to flee and decision distributions across six threat and reward conditions. Green dashed horizontal lines mark *x* = 0; blue dashed horizontal lines mark *x* = *x*_thr_; red dashed vertical lines mark the onset of each looming stimulus; blue vertical lines mark the median. (G) Heatmap of decision scores as a function of threat gain and reward value. White and red stars indicate fitted parameters for the early and late phases of the reward–threat paradigm, respectively; upward and downward pink triangles indicate fitted parameters for dominant and subordinate mice in the social-threat paradigm, respectively. (H) Heatmap of latencies to flee as a function of threat gain and reward value. (I) Vigilance as a function of input strength, illustrating how the indirect effect of reward on defensive decisions via vigilance depends on the baseline vigilance level. (J) Schematic showing how threat intensity, reward value, and vigilance jointly determine defensive decisions. Color saturation indicates defense likelihood.

In the second step, we identify threat gain and reward value across different threat and reward conditions in the late phase (Figures S9C–D). Consistent with experimental findings that decisions under low threat are primarily driven by reward value, the fitted reward-value parameter increases across reward conditions (0, 0.08, and 0.25 for no reward, water, and sucrose, respectively). With these parameters, the model reproduces the observed reduction in escape likelihood without changes in latency to flee (Figures 5A, 5C, and 5E). The same reward-value parameters are then used to model the decision-making process under high-threat conditions. Here, the fitted threat-gain parameter increases across reward conditions (1.44, 1.8, and 2.55 for no reward, water, and sucrose, respectively), counteracting—and ultimately reversing—the influence of reward value. The model again reproduces the experimentally observed increase in escape decisions and decrease in latency to flee (Figures 5B, 5D, and 5F).

To assess the model’s predictive power, we simulate how decision score (see Materials and Methods) and latency to flee vary as functions of reward value (*r*) and threat gain (*β*). Increasing reward generally reduces decision score and increases latency to flee, whereas increasing threat gain has the opposite effect (Figures 5G–H, and S9E–G). These relationships are confirmed by a simplified deterministic model (Figures S9H–K, see Materials and methods). Notably, reward can also indirectly promote escape via elevated vigilance, and the magnitude of this effect depends on the level of baseline vigilance, defined as vigilance in the absence of reward (Figure 5I). This dual influence explains the absence of a reward effect in the early phase under both threat conditions: high vigilance shifts the operating range to a region where reward’s indirect promoting effect is balanced by its direct suppressing effect on escape decisions.

We next apply this model to the social-threat paradigm. The behavior of dominant mice is captured by a combination of lower reward value and higher threat gain compared to subordinates (pink triangles in Figures 5G–H). This aligns with the experimental finding that dominant animals spent less time in the reward zone and fled more quickly (Figures 4O and 4J). Together, these results demonstrate that the model robustly captures key behavioral features across phases and paradigms, providing a unified mechanistic account of innate decision-making under threat (Figure 5J).

## 3 Discussion

When confronted by a predator, an animal must rapidly decide whether to escape, freeze, or continue its ongoing behavior. This life-or-death decision is especially critical for rodents facing aerial predators. Although mice are social animals and most predator encounters occur during foraging, it remains unclear how they integrate perceived threat and reward when making defensive decisions, or how these decisions are further influenced by social hierarchy. Using a foraging-based paradigm, we demonstrate that responses to visual threat are shaped by threat intensity, reward value, social rank, and internal state, and that this integrated decision process can be captured by a drift-diffusion leaky integrator.

### 3.1 Main findings

We designed a behavioral paradigm to simulate how mice respond to visual threats during foraging (Figure 1). Mice adapted rapidly to repeated looming stimuli, showing substantial inter-individual variability in the rate of habituation (Figure 2). While threat intensity robustly shaped behavior in both early and late phases of habituation, reward exerted a clear influence only during the late phase (Figure 3). Specifically, increasing threat intensity shifted behavior toward more defensive responses. Reward, in contrast, produced a context-dependent effect: under low-threat conditions, higher reward values suppressed defensive responses, consistent with value-based decision theory, whereas under high-threat conditions, they promoted escape decisions, likely through heightened vigilance. Innate decision-making was further shaped by social hierarchy (Figure 4): dominant mice exhibited a stronger bias toward defensive behaviors, whereas subordinates were more reward-driven and less likely to flee. Finally, we developed a drift-diffusion leaky integrator model that captures the key features of the observed behavioral patterns, providing a unified framework for understanding how threat, reward, and social context interact to shape innate defensive decisions (Figure 5).

### 3.2 Relation to earlier work

Aerial predators pose a significant threat to rodents. Previous studies have shown that overhead visual stimuli in the laboratory elicit robust defensive behaviors in rodents (Wallace et al., 2013; Yilmaz and Meister, 2013). Because these stimuli can mimic distinct predatory actions, the resulting behavioral responses depend on the physical properties of the stimulus. For example, an overhead expanding dark disc mimicking an approaching aerial predator preferentially triggers escape rather than freezing (Yilmaz and Meister, 2013), while a small black moving dot mimicking a cruising predator predominantly induces freezing behavior (De Franceschi et al., 2016). These findings suggest that the response is not a simple reflex but involves a decision-making process. Additional stimulus properties that influence action selection include contrast, speed, size, and shape (De Franceschi et al., 2016; Evans et al., 2018; Yang et al., 2020). Support for a decision-making process also comes from its modulation by environmental factors and experience: mice freeze more when no refuge is available (Wei et al., 2015) and quickly learn that the looming stimulus poses no real threat (Lenzi et al., 2022; Vale et al., 2017; Zhong et al., 2023). This innate decision-making process is observed across invertebrates and vertebrates (Evans et al., 2019).

Methodologically, the present study differs from earlier work in two aspects. First, instead of manually annotating behavioral responses, we employed a machine learning-based approach to classify behavioral decisions. Similar strategies have been used in previous studies of defensive behaviors (Fratzl et al., 2021). In our dataset, this approach substantially reduced the labor required for labeling and minimized misclassifications due to inter-individual variability. Second, unlike previous studies using 2D arenas, we designed a linear arena to simulate foraging conditions, where the nest and reward zone were separated by a long corridor. Importantly, the decision patterns observed in our paradigm are similar to those reported in 2D environments (Yang et al., 2020), and mice rapidly adapted to the looming stimulus after a few trials, consistent with previous findings (Lenzi et al., 2022).

One concern with the linear arena design was that the looming stimulus was usually presented at the end of the arena, raising the possibility that mice simply retreated upon reaching a physical boundary. To address this, we presented the stimulus between the foraging mice and the nest (Figure S2A). Under this condition, mice rapidly fled toward the nest—and thus toward the threat—rather than away from it (Figure S2B, Video 5). This behavior indicates that their escape responses are not simple reflexes, but instead incorporate the safety of the nest into their decisions.

Fanselow and Lester’s predatory imminence continuum theory posits that defensive behaviors are graded according to perceived threat intensity (Fanselow and Lester, 1988). To test whether this framework applies to aerial threats, we varied both prey–threat distance and prey–safety distance. As prey–threat distance increased, mice showed less direct escape behavior, with longer latencies to flee and slower escape speeds (Figures S2C–G, Videos 6–8), indicating that defensive responses scale with threat proximity. These effects cannot be explained by failure to detect the stimulus at longer distances; if that were the case, foraging speed would remain unaffected. Instead, mice slowed as they approached the threat in the 75-cm condition (Figure S2D). In contrast, prey–safety distance did not significantly influence defensive behaviors (Figures S2H–I). One interpretation is that once an aerial threat is detected, the urgency to escape overrides evaluation of refuge proximity. To further verify this, we introduced barriers that lengthened the return path to the nest. Even under these conditions, defensive behaviors remained unaffected by prey–safety distance (Figures S2J–K, Video 9).

### 3.3 Economic and social modulations of innate decision-making under threat

Value-based decision-making—where choices arise from comparisons of the subjective value of expected outcomes—is a well-established framework for understanding behavior across species (Rangel et al., 2008). Prospect theory, which accounts for decision-making under risk in humans (Kahneman and Tversky, 1979), has been applied to learned decision-making in rodents (Constantinople et al., 2019). Here, we demonstrate that value-based decision-making also extends to innate defensive behaviors: when confronted with a threat, mice integrate perceived reward and threat value to make their decisions.

Importantly, perceived value is determined not only by the physical properties of the stimulus but also by the animal’s internal state. In the resting state, for example, perceived stimulus strength scales with physical intensity according to the Weber–Fechner law. In our paradigm, perceived threat value is strongly modulated by vigilance, a state of heightened and sustained attention to potential danger. This vigilance-dependent modulation substantially alters how the same looming stimulus is evaluated and how decisions are made.

Consistent with value-based evaluation, increasing threat intensity reliably increased defensive responses across reward conditions and behavioral phases. These changes reflect differences in perceived threat rather than stimulus detection, as evidenced by coordinated changes in escape latency, distance, and speed, and further supported by trial-by-trial inspection confirming reliable stimulus detection across contrast levels (Figure S3B).

Reward-dependent behavior also follows value-based principles, but in a vigilance-dependent manner. In nature, foraging often occurs under predation risk, and evolutionary pressure has favored prey that maintain heightened vigilance in the presence of high-value rewards. Consistent with this, more valuable rewards increased both perceived reward value and—via elevated vigilance—perceived threat value. These opposing effects jointly shaped the escape decision.

In the early phase, baseline vigilance is already high because looming represents an innate threat. Under this high-vigilance condition, more valuable rewards elevates both perceived reward and threat value; these opposing effects counterbalance each other, resulting in no detectable behavioral differences across reward conditions. With repeated exposure, habituation to looming reduces vigilance, shifting the operating range leftward along a sigmoidal vigilance-intensity function (Figure 5I). This sigmoidal form is biologically plausible, as vigilance is bounded by a relaxed-state minimum and by cognitive capacity limits. This shift has opposite consequences under low-versus high-threat conditions. Under low threat in the late phase, reduced vigilance compresses the dynamic range, weakening the indirect effect of reward on perceived threat and allowing perceived reward value to dominate decisions. Under high threat, however, the vigilance function remains within a steeper region of the sigmoid, where increases in reward effectively elevate vigilance and thus perceived threat. As a result, reward produces the opposite behavioral effect under high threat. Together, these dynamics explain why reward effects are absent early but emerge after habituation in a threat-dependent manner.

Social rank-dependent differences in defensive behavior also fit within this framework. Even under identical conditions, dominant mice perceived higher threat values—driven by their elevated vigilance—along with lower reward values, consistent with their more risk-averse behavior. Similar status-dependent escape patterns have been reported across species, including voles (Kleiman et al., 2014) and crayfish (Krasne et al., 1997), suggesting an evolutionarily conserved strategy.

Because vigilance declines with habituation, factors that modulate the rate of habituation also influence defensive behavior. Although individuals vary, several patterns emerged. Mice habituated more rapidly under high-threat or high-reward conditions (Figure 2I). Dominant mice habituated more slowly than subordinates, consistent with their elevated vigilance (Figures 4H and S7A). Housing conditions also altered habituation: group-housed animals adapted more slowly to looming than single-housed animals, and dominant–subordinate pairs habituated even more slowly than group-housed mice (Figures 2I and S7A). These patterns indicate that stable social relationships sustain high vigilance and slow habituation—an evolutionarily conserved strategy that may enhance survival. Importantly, vigilance should not be conflated with stress or anxiety-like states. The slower habituation observed in group-housed mice reflects sustained sensitivity to repeated threat exposure rather than elevated anxiety. Thus, our data do not contradict the concept of social buffering; rather, they are consistent with it, as pair-housed mice exhibited reduced defensive responses compared with individually housed animals (Lenzi et al., 2022). Furthermore, in an independent study (Li et al., 2026), we observed that mice tested in the presence of a social partner displayed attenuated responses to looming stimuli relative to those tested alone. These findings suggest that social interactions can attenuate defensive responses while prolonging vigilance during repeated threat exposure.

One limitation of the present study is that our experiments were conducted exclusively in male mice to avoid the confounding effects of behavioral variability associated with the female estrous cycle. Extending this paradigm to females—with careful control for estrous cycle—will provide a more comprehensive understanding of how economic and social factors modulate defensive decision-making.

### 3.4 A proposed role for the superior colliculus in value integration

What neural circuits underlie the economic and social modulations of decision-making under threat? A growing body of research implicates the SC as a key node in looming-evoked defensive behaviors (Evans et al., 2018; Shang et al., 2015; Wei et al., 2015). In parallel, reward is encoded by dopaminergic neurons in the ventral tegmental area (VTA) (Cohen et al., 2012; Schultz, 1998) and serotonergic neurons in the dorsal raphe nucleus (Liu et al., 2014; Miyazaki et al., 2011), which project widely to regions such as the ventral striatum (Schultz et al., 1992), orbitofrontal cortex (Tremblay and Schultz, 2000), and cerebellum (Wagner et al., 2017). Furthermore, social status modulates behavior via circuits involving the medial prefrontal cortex (mPFC) (Kingsbury et al., 2019; Wang et al., 2011) and serotonergic signaling (Edwards and Kravitz, 1997; Raleigh et al., 1991; Yeh et al., 1997).

We propose the SC as a central integrative hub for mediating the economic and social modulation of defensive decision-making. First, neurons in the superficial (sSC) and intermediate (iSC) layers of the medial SC are critical in detecting looming threats and triggering defensive responses (Evans et al., 2018; Li et al., 2023, 2020; Wei et al., 2015). Second, two lines of evidence support reward modulation of the SC. On one hand, neurons in the medial iSC express dopamine and serotonin receptors (Mooney et al., 1996; Woolrych et al., 2021), with D1 receptors enriched in the sSC and D2 receptors in the iSC, suggesting that threat signals can be integrated with reward- and social-related information in a layer-dependent manner. Indeed, reward-related signals have been reported in the medial sSC (Baruchin et al., 2023). On the other hand, whereas the medial SC preferentially mediates defensive responses, the lateral SC promotes approach behaviors toward rewarding stimuli, such as food or prey (Comoli et al., 2012; Krauzlis et al., 2013), suggesting that medial–lateral interactions within the SC may play a critical role in resolving threat–reward trade-offs during decision-making. Third, social modulation can be conveyed through mPFC inputs and serotonergic pathways, providing a route for social status to directly bias SC computations underlying defensive decisions (Edwards and Kravitz, 1997; Kingsbury et al., 2019; Raleigh et al., 1991; Wang et al., 2011; Yeh et al., 1997).

Finally, SC computations are further shaped by contextual and experience-dependent signals that jointly influence economic and social modulation of behavior. The medial iSC receives inputs from the retrosplenial cortex (RSC) and hippocampus (Benavidez et al., 2021; Campagner et al., 2023), providing spatial and mnemonic context for evaluating threat and reward contingencies (Calvin et al., 2025). Consistently, looming stimuli engage hippocampal CA1 neurons (Conway et al., 2025). Moreover, repeated exposure induces habituation and fear extinction through a cortico-subcortical pathway in which higher visual areas recruit inhibitory neurons in the ventral lateral geniculate nucleus (vLGN) to suppress medial iSC activity (Fratzl et al., 2021; Mederos et al., 2025; Salay and Huberman, 2021), indicating experience-dependent reweighting of SC-mediated responses. These threat, reward, social, and contextual signals are integrated in the iSC and conveyed to the deep SC, which interfaces with downstream brainstem and midbrain effector systems, including the periaqueductal gray (PAG) and substantia nigra dopaminergic neurons (Comoli et al., 2003), enabling rapid transformation of integrated signals into defensive and approach actions.

While the anatomical and functional evidence support the SC as a central integrative hub, we emphasize that the neural architecture underlying such decisions is far more complex. The precise integration of internal state, social rank, and external threat likely emerges from the dynamic interplay of distributed circuits, including prefrontal, limbic, and neuromodulatory systems. Dissecting how these circuits interact to shape defensive decisions remains an important direction for future research.

### 3.5 Mathematical modeling

The proposed drift-diffusion leaky integrator model builds on an integrator model of internal state (Gibson et al., 2015) and extends it by incorporating a reward-driven drift-diffusion process (Ratcliff, 1978). Conceptually, the integrator component in our model aligns with Lorenz’s “hydraulic” model of motivation, which describes how internal drives shape behavior (Lorenz, 1950). While our model resembles the leaky integrator used to model escape behavior in flies (Gibson et al., 2015), it differs in several ways. First, the earlier model lacks a reward-related drift component. Second, while that model treats the looming effect as a delta function, our model allows the evidence level to vary continuously with stimulus size. Third, the influence of vigilance is absent in that model. A similar model has been proposed (Evans et al., 2018), but it did not incorporate reward and vigilance. Note that the evidence level for escape in our model is not equivalent to fear level (Anderson and Adolphs, 2014); rather, when it crosses a threshold, the animal may enter a fear state.

Below, we briefly discuss the roles of individual parameters. The leakage rate *α* represents a drive toward the resting state. Perceived threat is modeled as the product of the threat gain *β* and the sensory input *s*(*t*) and promotes escape decisions, whereas perceived reward *r* drives the decision away from escape. The threat gain increases not only with threat intensity but also with reward value via its influence on vigilance. The effect of reward on vigilance depends on the operating region of the sigmoidal vigilance function. This operating region is set by the baseline vigilance level (Figure 5I) and reflects habituation to repeated looming threats. Furthermore, threat gain and reward value have distinct effects on escape latency: higher threat gain exponentially shortened latency, whereas higher reward value increased it linearly (Figures S9F–G). These effects are confirmed using a simplified deterministic version of the model (Figure S9J–K; see Materials and Methods).

Lastly, the diffusion rate *δ* plays a critical role, particularly under low-threat conditions where the average evidence trajectory fails to reach the threshold. This parameter may reflect fluctuations of the animal’s internal state. The diffusion rate captures the trial-to-trial variability: even small amounts of noise can accumulate over time, resulting in large differences across individual trials and contributing to variability in escape latency. Overall, this computational framework not only accounts for our experimental observations but also offers a quantitative foundation for studying innate decision-making across species. Understanding how these parameters are implemented at the circuit level is an exciting direction for future research.

## 4 Materials and Methods

Key Resources Table.

| Reagent type (species) or resource | Designation | Source or reference | Identifiers |
| --- | --- | --- | --- |
| strain, strain background (*M. musculus*, male) | C57BL/6J | Chinese Institute for Brain Research, Beijing |  |
| chemical compound, drug | sucrose | Sigma-Aldrich | Cat#:V900116 |
| software, algorithm | Python v3.9 | Python Software Foundation | RRID:SCR_008394 |
| software, algorithm | R v4.2.1 | R Project for Statistical Computing | RRID:SCR_001905 |

### 4.1 Animals

Male C57BL/6J mice were group-housed on a 12-hour light/12-hour dark cycle and used at 2–3 months of age. Behavioral experiments were conducted during the light phase. All experimental procedures were performed under the animal welfare guidelines and approved by the Institutional Animal Care and Use Committee at the Chinese Institute for Brain Research, Beijing.

### 4.2 Behavioral platform

The behavioral platform consisted of a nest, a linear arena, a reward-delivery port, and an automated tracking system that included radio frequency identification (RFID) and real-time position detection. As illustrated in Figure 1A, the nest (40 (L) × 20 (W) × 30 (H) cm) and the linear arena (100 × 10 × 30 cm) were made of infrared-transmitting acrylic to allow unobtrusive behavioral monitoring. They were connected by a one-way tunnel (20 × 3 × 3 cm) for entering the arena and a 20 × 5 × 30 cm safe zone with a 5 × 5 cm one-way door for returning to the nest. Because it was not directly under the monitor, the safe zone was darker than the arena.

Mice were identified using implanted RFID tags, which were detected by an RFID reader positioned around the tunnel. Only one mouse was allowed to enter the arena at a time. Specifically, when all mice were in the nest, the door between the nest and the tunnel was open, while the door between the tunnel and the arena remained closed. When a single mouse entered the tunnel, as detected by the RFID system, the door to the nest closed, and the door to the arena opened, allowing the mouse to enter the arena. Mice received rewards at the end of the arena in the form of water or 10% sucrose. Licking time and reward volume were recorded for each trial.

To track mouse position in real time, an OpenMV camera with a 90° field-of-view lens was mounted on the ground, 110 cm beneath the arena. The mouse position was used to control the tunnel doors and to trigger stimulus presentation when the mouse entered the arena. A second camera (LBAS-U350-74M) with a lens (FA0615A) was also placed on the ground to record animal behavior at 30 frames per second.

### 4.3 Visual stimulation

Looming stimuli were presented on a 55-inch monitor (121 × 68 cm) suspended 32 cm above the arena. Visual stimuli were generated using PsychoPy in Python and aligned to the mouse’s real-time location. The stimulus consisted of a black disc that expanded ten times on a gray background (∼65 cd/cm^2^). On each expansion, the disc grew from 0° to 20° of visual angle at 40°/s, followed by a stationary phase lasting 0.3 s. The inter-stimulus interval was randomly varied between 1 and 2 minutes. Two stimulus contrasts (20% and 99%) were displayed by adjusting the luminance of the disc.

### 4.4 Economic modulation of innate decision-making

All mice were habituated to the behavioral platform for two days before the looming experiment. During both habituation and testing, three reward conditions were used: no reward, water, or 10% sucrose. Mice were not water-deprived in any of the groups. On the first day, five mice from the same home cage were placed in the nest for 30 minutes with all doors closed. Each mouse was then placed individually in the nest and allowed to explore the arena for 10 minutes under normal door operation; if a mouse entered the arena fewer than two times, the exploration period was extended until at least two arena visits were completed. Subsequently, all five mice were returned to the nest with all doors open and allowed to freely explore the arena for 2 hours to ensure that each mouse learned the reward location at the end of the linear arena. On the second day, each mouse was placed individually in the nest and given an additional 1 hour of exploration under normal door operation to further acclimate to the environment. The looming experiment was conducted the following day. An overhead looming stimulus of either low (20%) or high (99%) contrast was triggered when the mouse entered the trigger zone, defined as the region within 20 cm of the reward port. For each mouse, both the looming contrast and reward type remained constant across trials. In total, we collected and analyzed behavioral data from 590 trials across 62 mice.

To control for inter-individual variation in decision-making, we compared the same animal’s behavior under reward and no-reward conditions. Two groups of mice were used in this experiment. In the first group (*n* = 4), mice were first tested under the no-reward condition, followed by the reward condition. In the second group (*n* = 5), the order was reversed. For the reward condition, mice were water-deprived one day before exploration, and water was provided via the reward port. In the no-reward condition, mice were not water-deprived, and the reward port was removed. Before the looming experiment, mice were acclimated to the linear arena for two days as described above. On the third day, a looming stimulus with 20% contrast was displayed for five trials. In total, behavioral data from 84 trials across 9 mice were recorded and analyzed.

### 4.5 Social modulation of innate decision-making

To investigate the impact of social hierarchy on decision-making, we first determined the social rank of each mouse pair using the tube test (Lindzey et al., 1961; Wang et al., 2011), in which dominance is determined by the ability of one individual to force its opponent to retreat from a narrow tube. Mice were co-housed with a glass tube (3 cm diameter, 10 cm long) for one week, then trained to traverse a 30-cm tube 10 times per day for two consecutive days. On the third day, each pair was tested for up to six trials; if one mouse achieved four consecutive wins, it was designated the dominant individual; otherwise, the pair (1 out of 6 pairs) was excluded from further experiments.

Before the looming experiment, mice were water-deprived and allowed to explore the linear arena for two hours per day over three days, during which water rewards were delivered at the end of the arena. Over the following two days, the looming experiment was conducted with a stimulus contrast of 99% for two hours per day. After the looming experiment, mice continued exploring the arena for an additional two days, after which their social rank was reassessed with the tube test. All remaining pairs maintained a stable rank order. In total, behavioral data from 180 trials across 18 mice were recorded and analyzed.

To assess whether the tube test itself influenced defensive decision-making, additional looming experiments were conducted on four mouse pairs before and after the tube test. Specifically, the first looming experiment was conducted before the tube test (Figure S8A). Behavioral data from 72 trials across 8 mice were recorded and analyzed.

### 4.6 Behavioral quantification

Animal behaviors were quantified in three steps. First, two key points—the nose and tail base—were labeled and tracked in the recorded videos using DeepLabCut (Mathis et al., 2018). Tracked points with likelihood scores below 0.6 were excluded and replaced using linear interpolation. Locomotion speed was calculated for each frame and smoothed using a 0.3 s moving average window. Unless otherwise specified, tail base speed was used for subsequent analyses. Second, individual 30-second trials were extracted, each consisting of 10 s before, 8 s during, and 12 s after the looming stimulus.

Third, we defined 19 behavioral features related to locomotion speed, distance, and state transitions. Eleven features were related to speed and distance: (1) peak speed toward the reward port before stimulus onset; (2–4) maximum, mean, and coefficient of variation of speed after stimulus onset; (5) latency to peak speed; (6) hiding latency, defined as the time between stimulus onset and arrival at the safe zone (set to 20 s if the mouse did not reach the safe zone by the end of the trial); (7) escape distance, defined as the displacement from stimulus onset to offset; (8) distance to the nest at the end of the trial; (9) total distance traveled at speeds greater than 90% of the peak speed; (10) duration of movement at speeds greater than 90% of the peak speed; and (11) duration of time with both key points moving slower than 1 cm/s. Eight features were related to state transitions: (12) latency to flee, defined as the time from stimulus onset to the first escape episode, where an escape state was defined as movement exceeding 10 cm at a speed exceeding 10 cm/s; (13–15) average speed, distance traveled, and acceleration during the fastest escape episode; (16) latency to the fastest escape episode; (17) latency to the first stationary episode, where a stationary state was defined as both key points moving at less than 1 cm/s for more than 0.3 s; (18) total stationary duration and (19) longest stationary duration.

Finally, these features were fed into a random forest model with a maximum depth of 5 to classify behaviors. For trials in which neither escape nor stationary states were identified, latencies (features 12, 16, 17) were set to 20 s, distance (14) to 0 cm, speed (13) to 0 cm/s, acceleration (15) to 0 cm/s^2^, and duration (18, 19) to 0 s. The model was trained on 238 trials and tested on 102 trials, achieving an accuracy of 0.95 on the test set. Using this classifier, behavioral decisions in 3862 trials across 140 mice were categorized into four types: direct escape, escape after assessment, freezing, and no response (Figure S1). A decision score was calculated for each condition as *S* = 3*p_e_* + 2*p_ae_* + *p_f_*, where *p_e_*, *p_ae_*, *p_f_* denote the proportions of direct escape, escape after assessment, and freezing, respectively. To quantify how quickly animals recovered from a fear state, we measured recovery time. For escape trials, recovery time was defined as the hiding duration, measured from entry into the safe zone to re-entry into the arena. For freezing trials, recovery time was defined as the interval between the onset of the first freezing episode and the termination of the last freezing episode.

To quantify the influence of reward, the reward zone was defined as a region within 10 cm of the reward port, and the duration in the reward zone was defined as the time spent within this zone during the 20 s following stimulus onset. Vigilance was assessed using latency to flee, foraging interval (time between two consecutive entries into the trigger zone), and foraging speed (average speed while approaching the trigger zone before stimulus onset).

To segment the trials into two phases, we performed principal component analysis on a feature matrix containing decision score, escape distance, duration in the reward zone, peak escape speed, and latency to flee across the first 10 trials. We extracted the first principal component (Figures S3C–D), which captured the majority of the variance, and used its learning curve to detect transitions in behavior. Change points in the learning curve were detected using the ruptures method (Gallistel et al., 2004; Truong et al., 2020).

### 4.7 Behavioral modeling

Parameters in the drift-diffusion leaky integrator model were estimated in two steps by minimizing a loss function using a grid search. In the first step, we fit our model to behavioral data from the no-reward condition, setting *r* = 0 and *β* = 1 for the low-threat condition, and estimated *α*, *δ*, *x*_thr_, and *β* for the high-threat condition. In the second step, we estimated *β* and *r* for the water and sucrose conditions under both threat conditions. The loss function combines fitting errors in the distribution of behavioral decisions (*L*_dec_), the median latency to flee (*L*_med_), and the variability of latency (*L*_sd_):

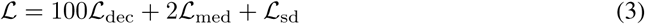

The decision-distribution mismatch is quantified using a mean cosine distance:

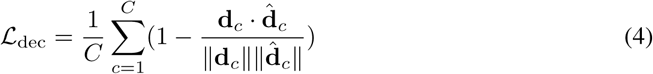

where **d***_c_* and **d^***_c_* denote the observed and model-predicted decision distributions for threat intensity *c*, and *C* = 2 is the number of threat conditions.

The median-latency loss is defined as

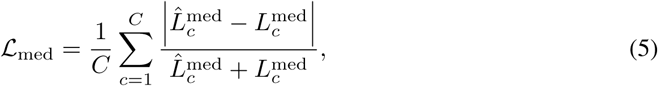

and the latency-variability loss is

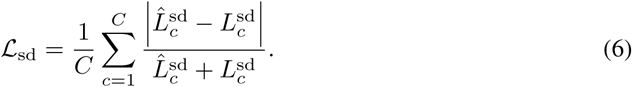

For the final fitted model (Equation 1), *α* = 1.78 and *δ* = 5.6. Under low-threat conditions, *β* = 1 for all reward conditions; under high-threat conditions, *β* = 1.44, 1.8, 2.55 for no reward, water, and sucrose, respectively. The reward values were *r* = 0, 0.08, 0.25 for no reward, water, and sucrose, respectively, under both threat conditions. In Equation 2, the decision threshold was *x*_thr_ = 0.71.

To further understand how threat and reward shape decisions, we analyzed a deterministic simplification of the model:

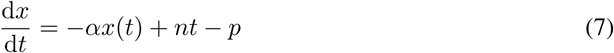

where *p >* 0 denotes the perceived reward value and *n >* 0 determines the rate at which threat-driven input increases over time.

With the initial condition *x*(0) = 0, the solution is

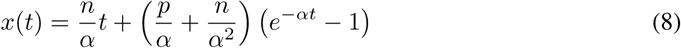

This solution reveals a competition between two components. First, a linear term driven by threat accumulation promotes escape over time. Second, an exponentially decaying component reflects the combined influence of reward and threat.

### 4.8 Quantification and statistical analysis

No statistical method was used to predetermine sample size. The Shapiro–Wilk test was applied to assess the normality of data distributions. For comparisons between two groups, a two-sample *t*-test or paired *t*-test was used for normally distributed data; otherwise, the Mann–Whitney *U* or Wilcoxon signed-rank test was applied. For multi-group comparisons, one-way or two-way analysis of variance (ANOVA) was used when data were normally distributed, followed by Tukey’s *post hoc* test. For non-parametric multi-group comparisons, the Kruskal–Wallis test (for one factor) or the Scheirer–Ray–Hare test (for two factors) was applied, followed by Dunn’s *post hoc* test with Holm correction. The chi-squared test was used to analyze categorical variables, while paired categorical comparisons were assessed using the Stuart–Maxwell test. Detailed statistical information for each experiment is provided in the Results section and figure legends.

## Acknowledgments

We thank Lei Zhang, Xueting Sun, Haojun Sang, Qun Zhang, Bing Zhao, and Zhuofan Li in the CIBR Instrumentation Core for their help in designing and building the linear arena. Ling-yun Li was supported by the Natural Science Foundation of Beijing Municipality (5244028), the National Natural Science Foundation of China (32471071), and the R&D Program of Beijing Municipal Education Commission (1240030201). Ya-tang Li was supported by the National Natural Science Foundation of China (32271060), the Natural Science Foundation of Beijing Municipality (IS23073), and the startup fund from CIBR.

## Author contributions

Ya-tang Li supervised the project; Ya-tang Li, Zhe Li, and Yidan Sun designed the experiments; Zhe Li collected all the data; Zhe Li, Jiahui Wang, and Jialin Li analyzed the data; Ya-tang Li carried out the neural modeling; Zhe Li and Ya-tang Li prepared figures; Ya-tang Li, Ling-yun Li, and Zhe Li wrote the manuscript.

## Data and code availability

Data and code are available in a public GitHub repository (https://github.com/YatangLiLab/ li-2025-economic-social-modulation).

## Supplemental information

**Figure S1:**
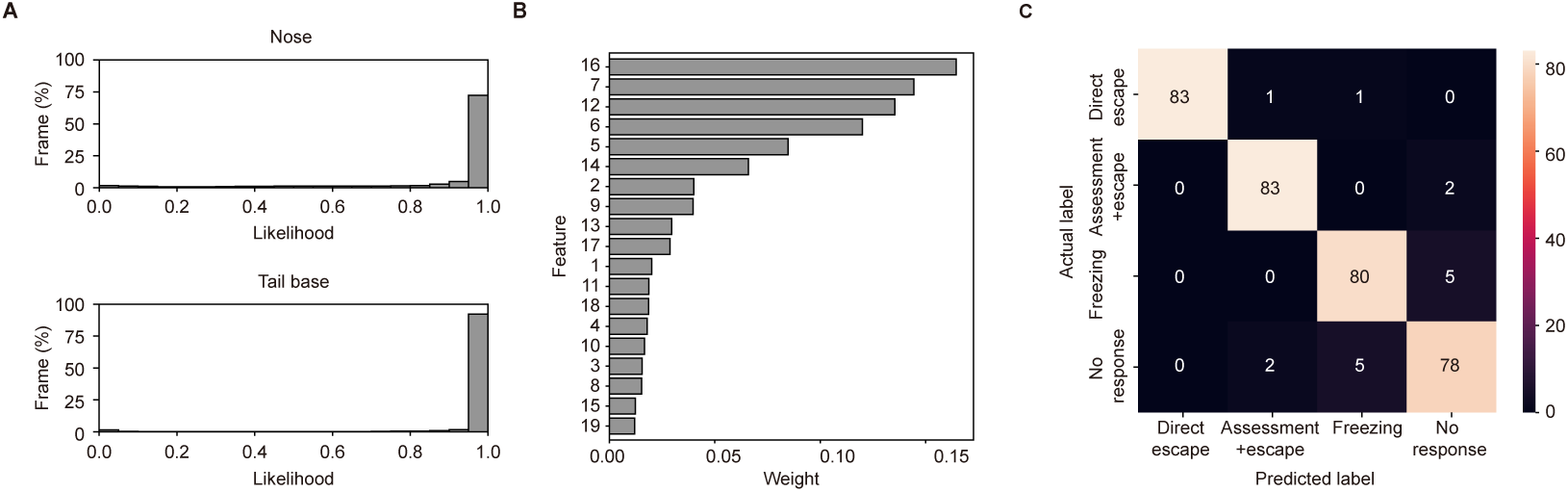
Model validation and feature analysis for automated behavioral classification. (A) Tracking accuracy of the mouse nose and tail base using DeepLabCut. (B) Feature weights in the random forest classifier. (C) Model performance evaluated by a confusion matrix on the test dataset.

**Figure S2:**
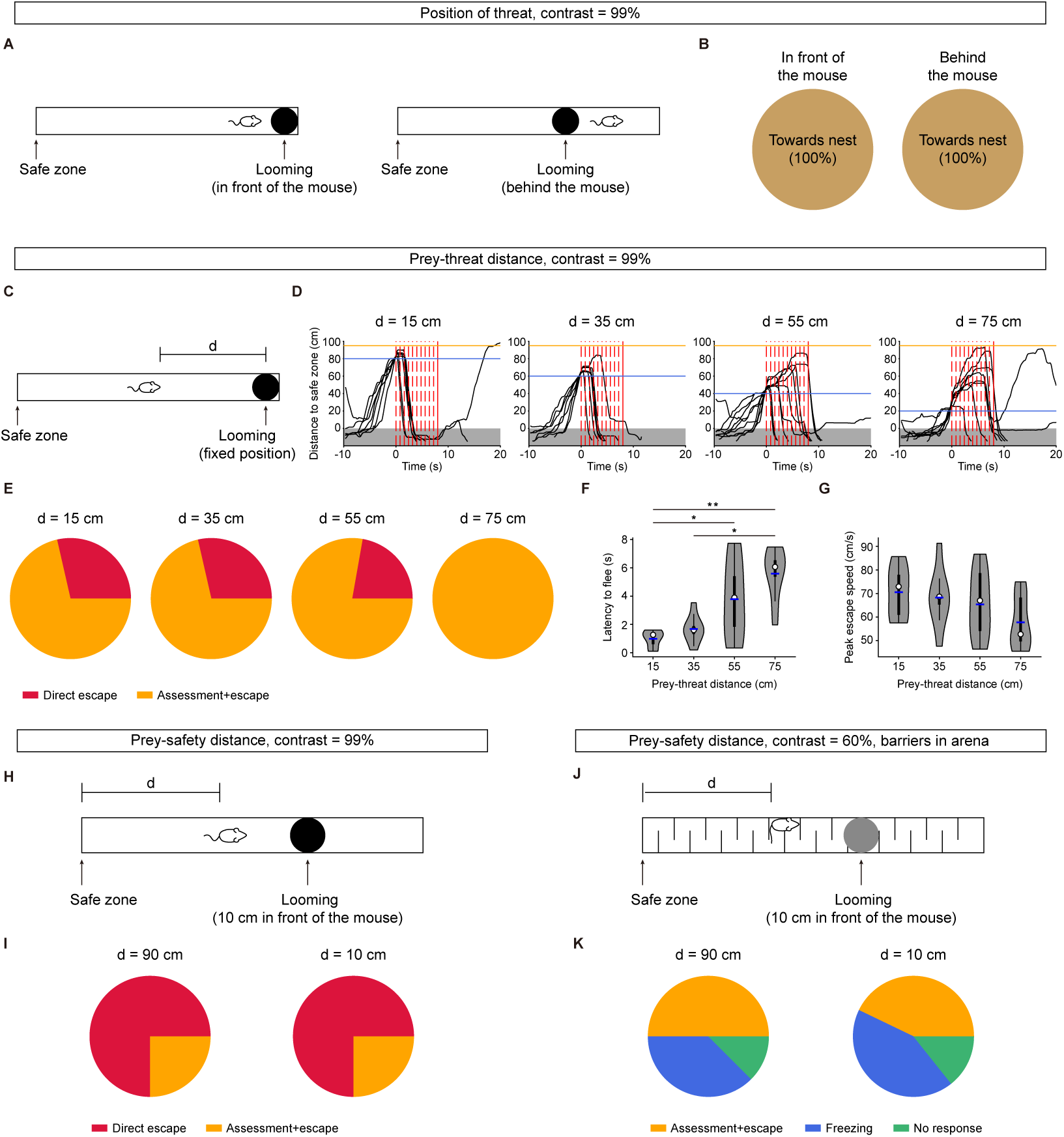
Effects of threat position, prey–threat distance, and prey–safety distance on defensive responses to looming stimuli. (A) Schematic of the experiment in which looming stimuli were presented either in front of or behind the mice during foraging. (B) Distributions of escape directions for front and rear stimulus positions. *n* = 97, 16 trials. (C) Schematic of the experiment in which looming stimuli were presented at the end of the linear arena with varying distances between the mouse and the threat. (D) Distance to the safe zone over time for four prey–threat distances. Red dashed lines mark the onset of each stimulus repetition; solid lines mark the end of the last repetition. Gray shading indicates the safe zone. Blue lines mark the mouse position at stimulus onset; orange lines mark the stimulus location. *n* = 7 (15 cm), 7 (35 cm), 9 (55 cm), and 8 (75 cm) trials from 5 mice. (E) Distribution of decisions across prey–threat distances. (F–G) Latency to flee and peak escape speed across prey–threat distances. Kruskal–Wallis test with *post hoc* Dunn’s test (Holm correction). (H) Schematic of the experiment in which looming stimuli were presented in front of the mouse at varying distances between the mouse and the safe zone. (I) Distribution of decisions across prey–safety distances. *n* = 4, 4 trials. (J) Schematic of the experiment in which low-contrast looming stimuli were presented at varying distances between the mouse and the safe zone in the linear arena with barriers. (K) Distribution of decisions across prey–safety distances in the barrier condition. *n* = 8, 7 trials. \**P <* 0.05, \*\**P <* 0.01.

**Figure S3:**
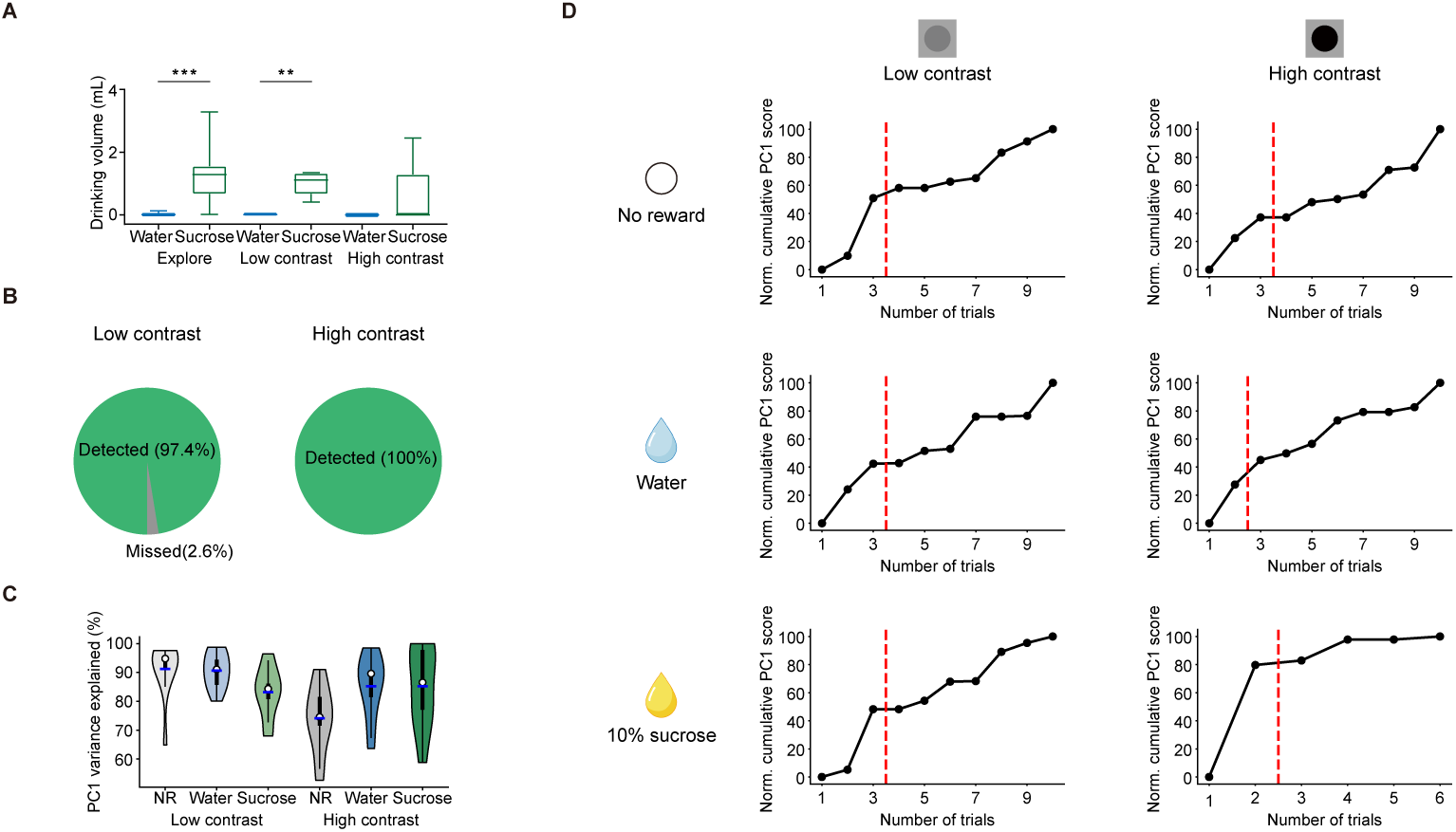
(A) Water and sucrose consumption during exploration and looming experiments under low- and high-contrast conditions. *n* = 10, 10, 5, 5, 5, 5 sessions; Mann–Whitney *U* test. Boxes represent the interquartile range (IQR), and whiskers show the full data range. (B) Pie chart showing the proportion of looming stimuli detected by mice in no-response trials under low- and high-threat conditions. Low contrast: *n* = 117 trials from 29 mice; high contrast: *n* = 2 trials from 2 mice. (C) Proportion of behavioral variance explained by principal component 1 (PC1) for individual mice across all conditions. *n* = 11, 13, 10, 10, 8, 10 mice. (D) Cumulative PC1 score for an example mouse in each condition. Red dashed lines mark the transition from the early to the late phase. \**P <* 0.05, \*\**P <* 0.01.

**Figure S4:**
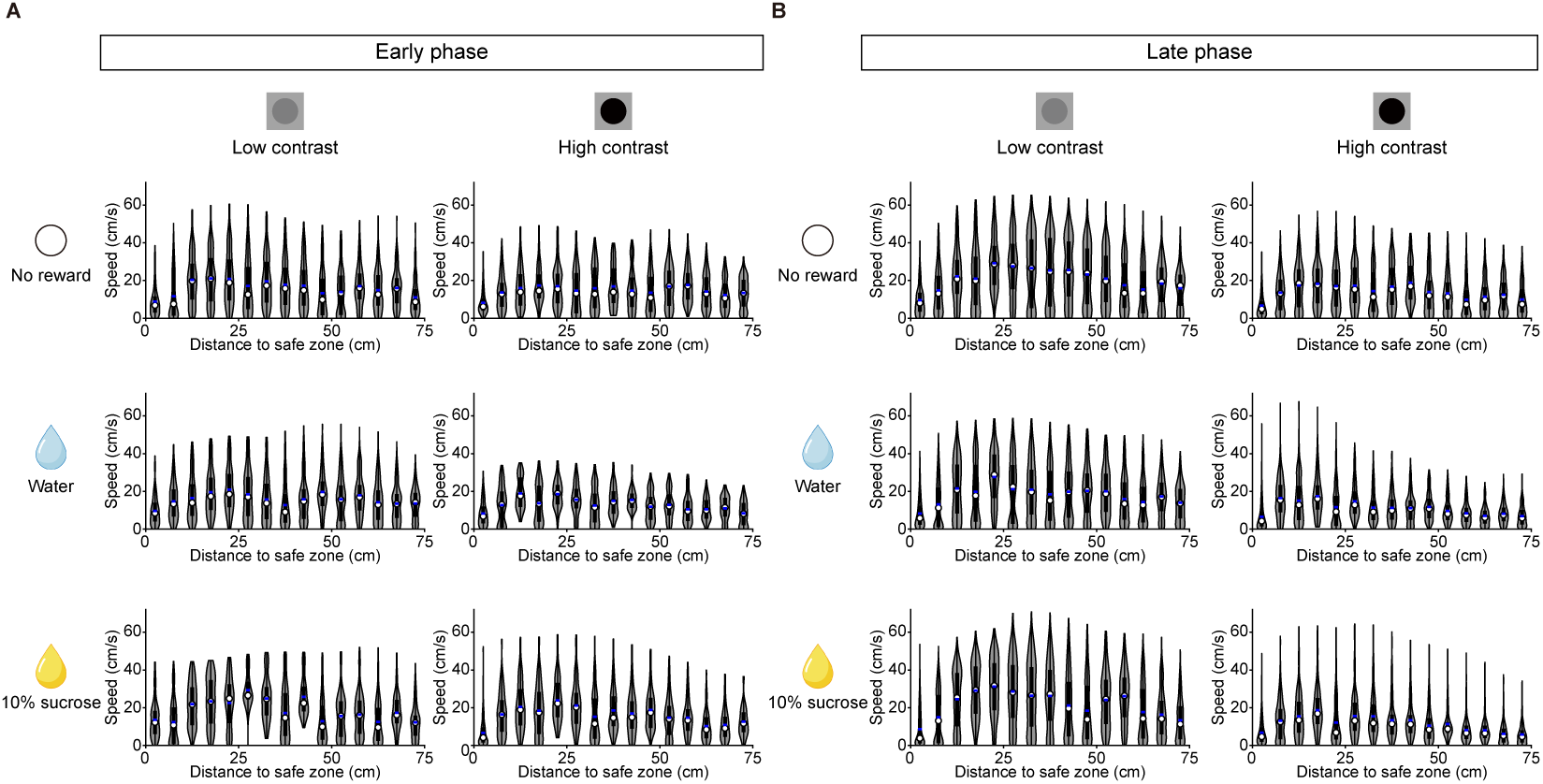
Distribution of foraging speed. (A) Distribution of foraging speed as a function of distance from the safe zone in the early phase across different threat and reward conditions. (B) Distribution of foraging speed in the late phase. Data are from 11 mice (no reward, low contrast), 13 mice (water, low contrast), 10 mice (sucrose, low contrast), 10 mice (no reward, high contrast), 8 mice (water, high contrast), and 10 mice (sucrose, high contrast).

**Figure S5:**
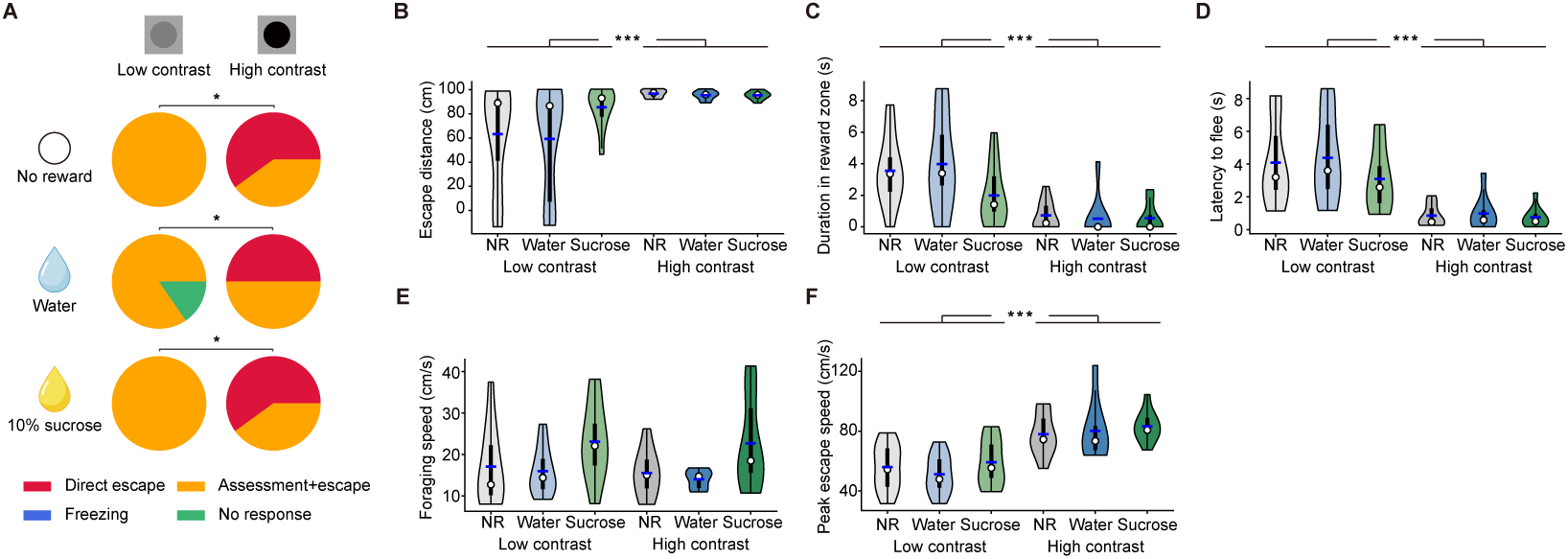
Behavioral responses to looming stimuli across threat and reward conditions in the first trial. (A) Distribution of decisions under six conditions. *n* = 11 (no reward, low contrast), 13 (water, low contrast), 10 (sucrose, low contrast), 10 (no reward, high contrast), 8 (water, high contrast), 10 (sucrose, high contrast) mice; chi-squared test. (B–F) Escape distance under threat, duration in the reward zone, latency to flee, foraging speed, and peak escape speed across conditions. *n* = 11, 13, 10, 10, 8, 10 mice for B, C, E, F; *n* = 11, 11, 10, 10, 8, 10 mice for D; Scheirer–Ray–Hare test with *post hoc* Dunn’s test (Holm correction). \**P <* 0.05, \*\*\**P <* 0.001.

**Figure S6:**
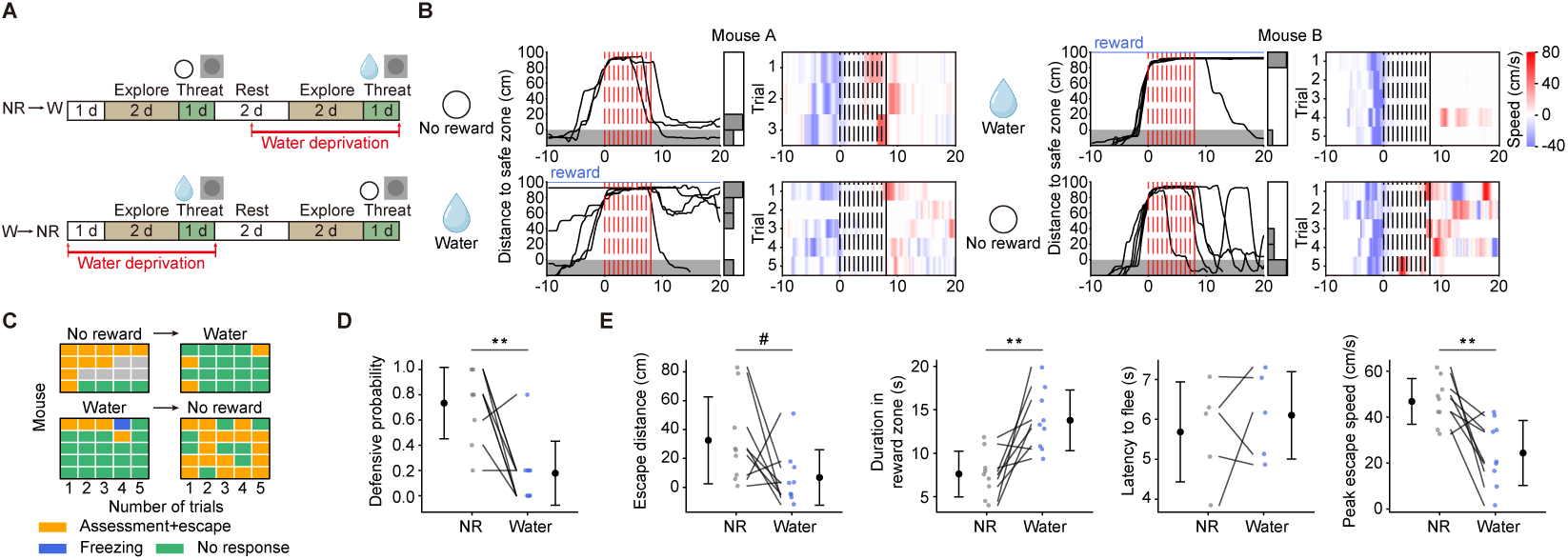
Reward modulation of defensive behavior to looming stimuli within the same mouse. (A) Experimental timeline illustrating how water reward influences innate decision-making within the same animal. (B) Distance to the safe zone and locomotion speed toward the safe zone across trials in response to low-contrast looming stimuli with and without water reward in two example mice. Left: no-reward condition followed by water-reward condition; right: water-reward condition followed by no-reward condition. (C) Decision patterns of nine mice across two sessions with 5 trials for each. Gray squares indicate trials where the mouse did not enter the arena within 30 minutes. (D) Defensive probability in no-reward and water-reward conditions. *n* = 9 mice; paired *t*-test. (E) Escape distance under threat, duration in the reward zone, latency to flee, and peak escape speed in no-reward and water-reward conditions. *n* = 5 mice for latency to flee and 9 mice for other measures; paired *t*-test. #*P <* 0.1, \**P <* 0.05, \*\**P <* 0.01.

**Figure S7:**
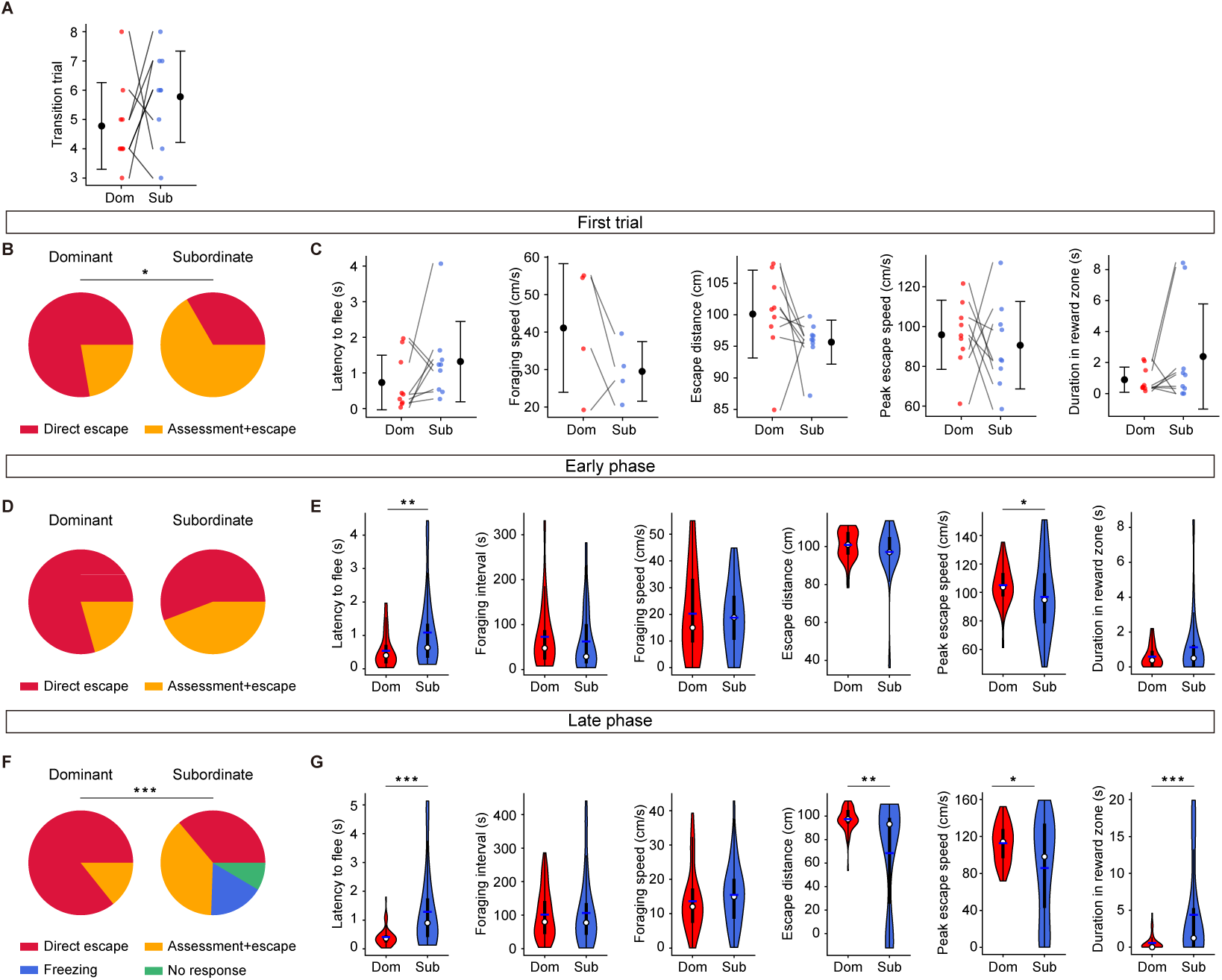
Behavioral responses to looming stimuli for dominant and subordinate mice in different phases. (A) Transition trial marking the start of the late phase for dominant and subordinate mice. *n* = 9 pairs. (B) Behavioral decisions for dominant and subordinate mice during their first threat exposure. *n* = 9 pairs; Stuart-Maxwell test. (C) Latency to flee, foraging speed, escape distance under threat, peak escape speed, and duration in the reward zone during the first threat exposure. *n* = 4 pairs for foraging speed; *n* = 9 pairs for other features; paired *t*-test. (D) Behavioral decisions for dominant and subordinate mice in the early phase. *n* = 34, 43 trials, respectively; chi-squared test. (E) Violin plots showing latency to flee, average foraging interval, foraging speed, escape distance under threat, peak escape speed, and duration in the reward zone for dominant and subordinate mice in the early phase. *n* = 34, 43 trials, respectively; Mann–Whitney *U* test. (F) Behavioral decisions for dominant and subordinate mice in the late phase. *n* = 56, 47 trials, respectively; chi-squared test. (G) Violin plots showing latency to flee, average foraging interval, foraging speed, escape distance under threat, peak escape speed, and duration in the reward zone for dominant and subordinate mice in the late phase. *n* = 56, 35 trials for latency to flee; *n* = 56, 47 trials for other measures. Mann–Whitney *U* test. For all panels: \**P <* 0.05, \*\**P <* 0.01, \*\*\**P <* 0.001.

**Figure S8:**
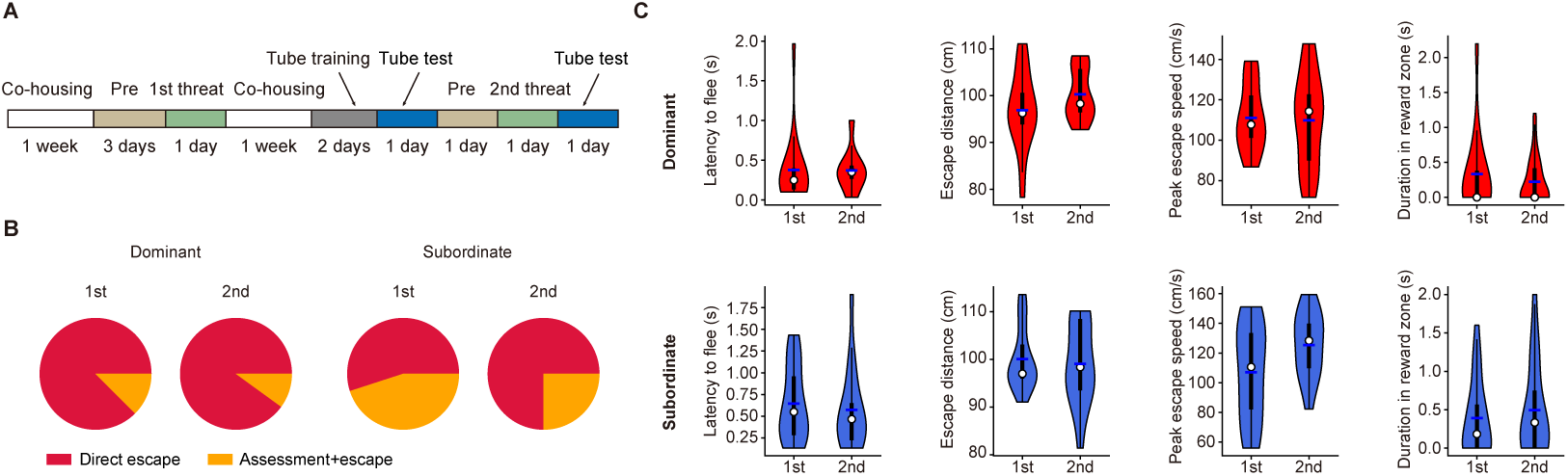
Comparison of behavioral responses to looming stimuli before and after the tube test. (A) Schematic timeline of the looming experiments before (first threat exposure) and after (second threat exposure) the tube test. (B) Behavioral decisions for dominant and subordinate mice during the first and second threat exposures. Dominant: *n* = 16 (first) and 20 (second) trials from 4 mice; subordinate: *n* = 20 (first) and 16 (second) trials from 4 mice; chi-squared test. (C) Violin plots showing latency to flee, escape distance under threat, peak escape speed, and duration in the reward zone for dominant (top) and subordinate (bottom) mice during the first and second threat exposures. *n* as in (B); Mann–Whitney *U* test.

**Figure S9:**
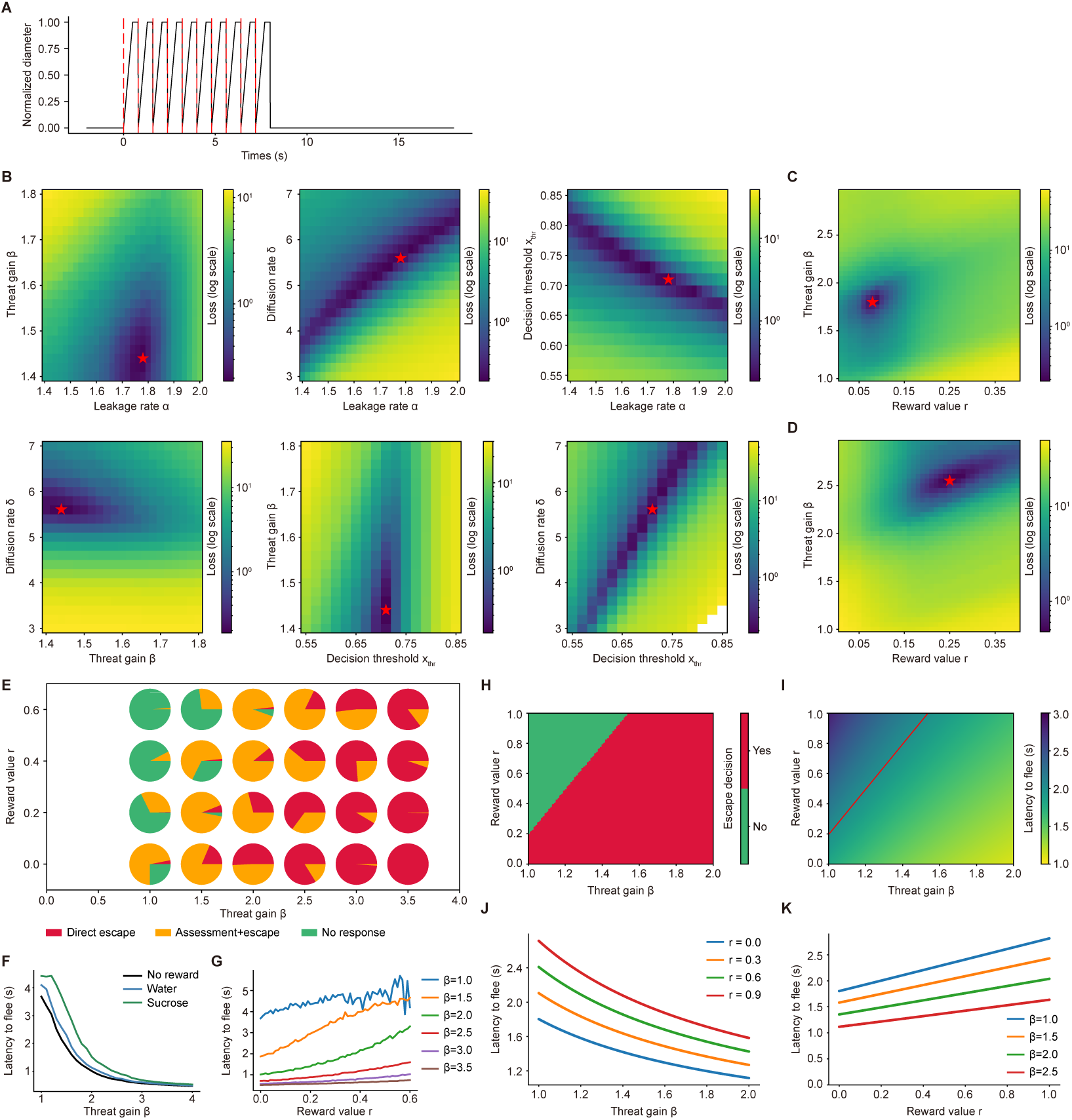
Fitting loss in the drift-diffusion leaky integrator model. (A) Temporal profile of the normalized looming stimulus diameter. (B) Loss landscapes for estimating the leakage rate, high-threat gain, diffusion rate, and decision threshold during the first-stage fitting. Red stars indicate the optimal parameter estimates based on experimental data in the late phase of the reward-related paradigm under the no-reward condition. (C) Loss landscapes for estimating the reward value and high-threat gain under the water-reward condition. Red stars mark the optimal fits to experimental data in the late phase of the reward–threat paradigm. (D) Same as (C), for the sucrose-reward condition. (E) Model-predicted distribution of decisions across threat gain and reward values. (F) Predicted latency to flee as a function of threat gain for varying reward values. (G) Predicted latency to flee as a function of reward value for varying threat gain. (H–I) Escape decisions and latencies to flee across threat gain and reward value predicted by a simplified deterministic model. (J–K) Same as (F–G), but for the simplified deterministic model.

## Video titles and legends

**Videos 1–4:** Example videos showing the four types of behavioral decisions in response to looming stimuli: direct escape, escape after assessment, freezing, and no response.

**Video 5:** Example video showing defensive responses to looming stimuli presented behind the mouse.

**Videos 6–8:** Example videos showing defensive responses at different prey–threat distances: 35 cm, 55 cm, and 75 cm.

**Video 9:** Example video showing defensive responses to looming stimuli in the linear arena with barriers.

